# Selective activation of estrogen receptors α and β: Implications for depressive-like phenotypes in female mice exposed to chronic unpredictable stress

**DOI:** 10.1101/758862

**Authors:** Rand S. Eid, Stephanie E. Lieblich, Paula Duarte-Guterman, Jessica A. Chaiton, Amanda G. Mah, Sarah J. Wong, Yanhua Wen, Liisa A.M. Galea

## Abstract

The estrogen receptor (ER) mechanisms by which 17β-estradiol influences depressive-like behaviour have primarily been investigated acutely and not within an animal model of depression. Therefore, the current study aimed to dissect the contribution of ERα and ERβ to the effects of 17β-estradiol under non-stress and chronic stress conditions. Ovariectomized (OVX) or sham-operated mice were treated chronically (47 days) with 17β-estradiol (E2), the ERβ agonist diarylpropionitrile (DPN), the ERα agonist propylpyrazole-triol (PPT), or vehicle. On day 15 of treatment, mice from each group were assigned to Chronic Unpredictable Stress (CUS; 28 days) or non-CUS conditions. Mice were assessed for anxiety- and depressive-like behaviour and hypothalamic-pituitary-adrenal (HPA) axis function. Cytokine and chemokine levels, and postsynaptic density protein 95 were measured in the hippocampus and frontal cortex, and adult hippocampal neurogenesis was assessed. Overall, the effects of CUS were more robust that those of estrogenic treatments, as seen by increased immobility in the tail suspension test (TST), reduced PSD-95 expression, reduced neurogenesis in the ventral hippocampus, and HPA axis negative feedback dysregulation. However, we also observe CUS-dependent and -independent effects of ovarian status and estrogenic treatments. The effects of CUS on PSD-95 expression, the cytokine milieu, and in TST were largely driven by PPT and DPN, indicating that these treatments were not protective. Independent of CUS, estradiol increased neurogenesis in the dorsal hippocampus, blunted the corticosterone response to an acute stressor, but increased anxiety-like behaviour. These findings provide insights into the complexities of estrogen signaling in modulating depressive-like phenotypes under non-stress and chronic stress conditions.

## Introduction

Major depressive disorder (MDD) is a stress-related psychiatric disorder that affects approximately twice more women than men (Kessler and Bromet, 2013; Salk et al., 2017). Given this disparity in prevalence rates, a focus on female-specific factors that may contribute to depression is warranted. Sex differences in prevalence rates first emerge around puberty (Salk et al., 2017), suggesting that ovarian hormones may increase risk for depression in women. However, periods associated with declines in ovarian hormones, including the postpartum and perimenopause, are associated with a heightened risk for depression in women (Cohen et al., 2006; Hendrick et al., 1998; Soares, 2014), supporting the notion that ovarian hormones may afford resilience, rather than risk. The complex roles of ovarian hormones in depression may be partly attributed to the diversity of estrogen signaling, which can be loosely categorized into genomic and non-genomic mechanisms (Björnström and Sjöberg, 2005).

Evidence from animal models suggests that 17β-estradiol may interact with classical estrogen receptors (ERs) to regulate affective function. However, this is primarily derived from studies that have used tests of depressive- and anxiety-like behaviour in the absence of any manipulations intended to induce depressive- or anxiety-like phenotypes (e.g. chronic stress). For example, the administration of 17β-estradiol or an ERβ but not ERα agonist to ovariectomized rats decreases anxiety- and depressive-like behaviour (Walf et al., 2004; Walf and Frye, 2005; Weiser et al., 2009; Yang et al., 2014). Moreover, the anxiolytic and antidepressant-like effects of 17β-estradiol are not observed in ERβ knockout mice (Rocha et al., 2005). Collectively these findings indicate that 17β-estradiol’s effects on affective behaviours may be primarily mediated via interactions with ERβ. Importantly, the effects ER activation in “healthy” animals may not mirror those in animals displaying depressive-like pathology, and as such although informative, these findings are certainly not conclusive. Indeed, a more recent study found that overexpression of ERα in the nucleus accumbens afforded resilience to the depressive-like outcomes after stress exposure in female mice (Lorsch et al., 2018), but the authors did not test the effects of ERβ overexpression. Thus, to date, the mediatory roles of ERα and ERβ have not been investigated systematically *and* within an animal model of depression. The current study therefore utilized the chronic unpredictable stress (CUS) model in female mice with the aim of dissecting the contribution of ERα and ERβ to the estrogenic modulation of depressive-like pathology.

Endocrine and immune systems engage in extensive crosstalk (Bereshchenko et al., 2018; Grossman, 1985; Olsen and Kovacs, 1996). 17β-Estradiol in particular possesses immunomodulatory properties that are mediated via interactions with both ERα and ERβ (Baker et al., 2004; Kovats, 2015; Saijo et al., 2011). For example, several *in vitro* and *in vivo* studies in females reveal the capacity for 17β-estradiol and selective ER modulators (SERMs) to ameliorate inflammation and microglial activation in response to inflammatory challenges (Baker et al., 2004; Brown et al., 2010; Ishihara et al., 2015; Vegeto et al., 2008, 2001). The anti-inflammatory properties of estradiol may be of particular interest within the context of depression, as inflammation in MDD has been well-documented and corroborated by animal models of stress exposure (Hodes et al., 2015; Miller and Raison, 2016). Indeed, meta-analyses show increased proinflammatory cytokines including IL-6 and TNF-α in blood, cerebrospinal fluid, and brains of individuals with MDD (Dowlati et al., 2010; Enache et al., 2019; Haapakoski et al., 2015). Despite this, there has been insufficient consideration of how sex and sex hormones can influence the association between depression and inflammation (reviewed in Eid et al., 2019b). It is plausible that the immunomodulatory effects of 17β-estradiol are implicated in depression and the outcomes of stress exposure, yet the receptor mechanisms by which 17β-estradiol can influence chronic stress-induced inflammation remain unclear.

The hippocampus retains the ability to produce new neurons in adulthood in a variety of species including humans (Kempermann et al., 2018; Moreno-Jiménez et al., 2019; Tobin et al., 2019). Adult hippocampal neurogenesis is reduced by chronic stress, promotes stress resilience, and is implicated in affective function and hypothalamic-pituitary-adrenal (HPA) axis regulation (Anacker et al., 2018; Schoenfeld and Gould, 2012; Surget et al., 2011), although most of this evidence is derived from male rodents. Importantly, 17β-estradiol influences adult hippocampal neurogenesis in females (reviewed in Mahmoud et al., 2016b) and its pro-proliferative effects are mediated via actions on both ERα and ERβ, at least acutely (Mazzucco et al., 2006). However, whether chronic activation of ERα and ERβ influences the survival of newly produced hippocampal neurons under basal or chronic stress conditions has not been investigated. Beyond hippocampal neurogenesis, both estrogens and stress exposure affect synaptic plasticity in a variety of brain regions, including the hippocampus and frontal cortex (Brinton, 2009; Li et al., 2004; McEwen, 2013; McEwen et al., 2016; Srivastava and Penzes, 2011). Estradiol and ERα or β agonist upregulate the expression of synaptic proteins in female rats (Waters et al., 2009), including the postsynaptic density protein 95 (PSD-95), a scaffolding protein which organizes synaptic elements that are important for synaptic function and plasticity (Kim and Sheng, 2004). Importantly, PSD-95 is reduced in the prefrontal cortex (PFC) of individuals with MDD (Feyissa et al., 2009) and in animal models of stress exposure (Kallarackal et al., 2013; Kim and Leem, 2016; Pacheco et al., 2017). Yet, whether and by what receptor mechanisms can 17β-estradiol influence the effects of stress on PSD-95 remains to be elucidated.

Dysregulation of the HPA axis is observed in a subset of individuals with MDD, and includes hypercortisolemia, aberrations in the circadian rhythm of cortisol, and impairments in HPA axis negative feedback function (Ising et al., 2007; Stetler and Miller, 2011). Importantly, the HPA axis interacts bidirectionally with the hypothalamic pituitary gonadal (HPG) axis (Reviewed in Goel et al., 2014). HPA-HPG interactions in females have been primarily investigated at baseline or in response to acute stressors; in general, 17β-estradiol stimulates HPA axis activity and impairs its negative feedback function (Goel et al., 2014), with opposing roles of ERα and ERβ (Weiser et al., 2010; Weiser and Handa, 2009). Importantly, these effects of 17β-estradiol may not translate to conditions of chronic stress exposure. Indeed, we have shown that long-term ovarian hormone deprivation in rats increased HPA axis negative feedback impairment under chronic stress exposure (Mahmoud et al., 2016), which parallels findings in MDD indicating greater negative feedback impairment in women post-menopause (Roy et al., 1986; Young et al., 1993). Therefore, we investigated the roles of ERα and ERβ in mediating the effects of estradiol on HPA axis function under chronic stress conditions.

In this study, we sought to determine whether 17β-estradiol conferred resilience in the face of CUS in adult female mice. We further employed a pharmacological approach to disentangle the contribution of ERα and ERβ to the effects of 17β-estradiol under non-stress and chronic stress conditions. We examined stress susceptible behaviours, markers of plasticity in the hippocampus and frontal cortex, central immune mediators, and HPA axis function. We hypothesized that ovariectomy would increase depressive-like endophenotype under non-stress conditions and further exacerbate the depressive-like outcomes of chronic stress exposure. Additionally, we expected estradiol treatment to rescue the effects of ovariectomy and that its actions would be differentially facilitated by ERα and ERβ.

## Methods

### Animals

Young adult female C57BL6/J mice (Charles River Laboratories; Quebec, Canada) arrived at our facility at approximately 8 weeks of age. All mice were group housed and given *ad libitum* access to food and water. Mice were maintained on a 12-hour light/dark cycle (lights on at 7:00am) in temperature- and humidity-controlled colony rooms (21LJ±LJ1°C and 50LJ±LJ10%, respectively). All procedures were conducted in accordance with the ethical guidelines of the Canadian Council on Animal Care and were approved by the Animal Care Committee at the University of British Columbia.

### Surgery

All mice received bilateral ovariectomy (OVX) or sham surgery at approximately 3 months of age, adapted from our previously described procedure in rats (Barha et al., 2015; Galea et al., 2018). Briefly, surgery was performed through two lateral skin and muscle incisions under isoflurane anesthesia (5% induction, 1-1.5% maintenance), with ketamine (50mg/kg; intraperitonially (i.p.)), xylazine (5mg/kg; i.p.), bupivacaine (2mg/kg; local), and Lactated Ringer’s Solution (10ml/kg; subcutaneously (s.c.)). Mice received a nonsteroidal anti-inflammatory analgesic at the time of surgery, and at 24- and 48-hours post-surgery (Anafen; 5mg/kg; s.c.) and recovered for 7-9 days prior to hormone/agonist treatment (see experimental timeline in **Fig. 1**).

**Figure 1.**
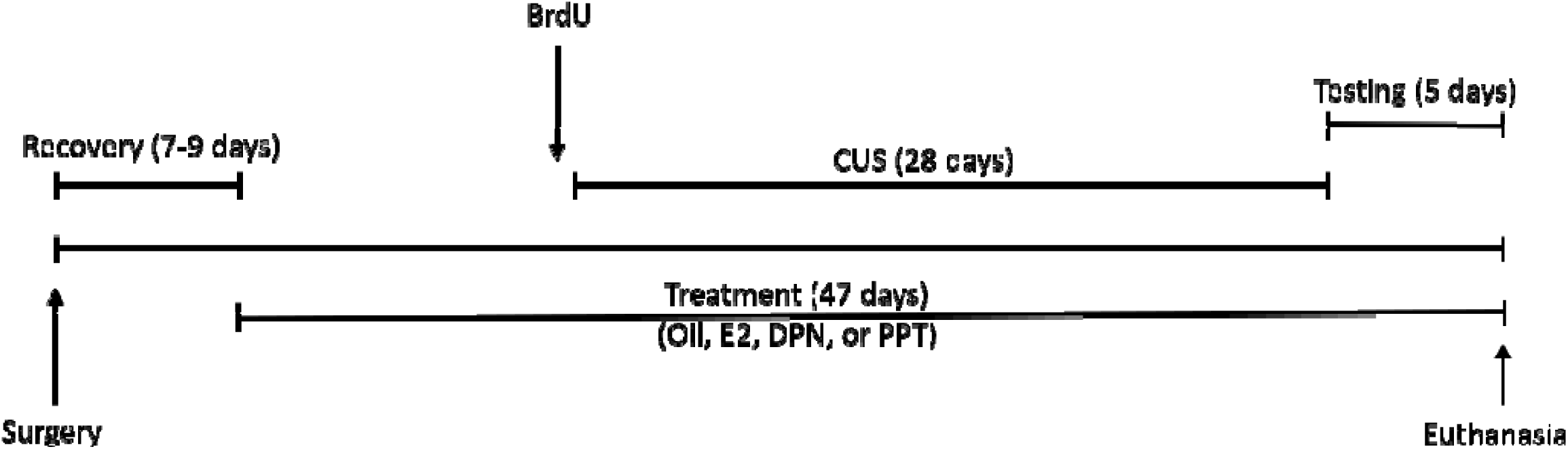
Timeline depicting the sequence of experimental events. Ovariectomy or Sham surgery was performed at approximately 3 months of age. Following 7-9 days of recovery, all mice received daily subcutaneous treatment (oil, E2, DPN, or PPT) until experimental endpoint (47 days total). On treatment day 14 all mice received two BrdU injections. Starting on treatment day 15, half the mice from each treatment group were subjected to chronic unpredictable stress (CUS; 28 days). Behavioural tests were administered starting the day after CUS in the following order (one test/day): forced swim test, sucrose preference test, novelty suppressed feeding test, and tail suspension test. On the final day of the experiment mice were subjected to the dexamethasone suppression test or the HPA axis stress reactivity test before tissue harvesting.

### Hormone and agonist treatment

Ovariectomized mice were randomly assigned to 1 of 4 treatment groups (n=26-28 per group), receiving daily s.c. injections of one of the following: (i) 17β-estradiol (E2; 0.04 mg/kg), (ii) diarylpropionitrile (DPN; 0.1 mg/kg; ERβ-selective agonist), (iii) propylpyrazole-triol (PPT; 0.1 mg/kg; ERα-selective agonist), or (iv) vehicle (sesame oil). To control for injections, sham-operated mice (n=20) also received daily s.c. vehicle injections (sesame oil). We therefore obtained the following 5 treatment groups: E2-treated OVX (E2), DPN-treated OVX (DPN), PPT-treated OVX (PPT), oil-treated OVX (OVX), and oil-treated sham-operated mice (Sham). DPN was chosen as it has a 70-fold greater affinity for ERβ over ERα (Meyers et al., 2001), and PPT was chosen as it has a 410-fold greater affinity for ERα over ERβ (Stauffer et al., 2000). 17β-estradiol has an equally high affinity to both ERα and ERβ, and its dose was chosen based on previous work and reports showing physiological circulating estradiol concentrations (Ciana et al., 2003; Harburger et al., 2007). Agonist doses were based on reports of behavioural efficacy (Clipperton et al., 2008; Walf et al., 2008). Treatment persisted for 47 days (see **Fig. 1**), beginning 14 days prior to CUS exposure. The choice to begin treatment prior to CUS was to examine the potentially protective, rather than therapeutic effects of 17β-estradiol or agonists.

### Bromodeoxyuridine administration

To label dividing progenitor cells and their progeny, 2 injections of bromodeoxyuridine (BrdU; 200mg/kg; i.p; 8 hours apart) were administered to all mice two weeks after hormone treatment had begun, and one day prior to the initiation of CUS (**Fig. 1**). This timing of BrdU administration was intended for the examination of the effects of CUS on the survival of new neurons, independent of effects on cell proliferation.

### Chronic Unpredictable Stress

Mice from each treatment group (Sham, OVX, E2, DPN, and PPT) were randomly assigned to chronic unpredictable stress (CUS) or no-CUS conditions, resulting in 10 groups overall (n = 10-14). CUS consisted of exposure to 13 stressors, applied twice daily in a pseudo-random order for a duration of 4 weeks (**Fig. 1**). The CUS protocol was adapted from published reports (Kreisel et al., 2014; R. Mahmoud et al., 2016; Wainwright et al., 2016) and stressors used are detailed in **Table 1**. Chronic unpredictable stress was chosen as it shows face, construct and predictive validity as an animal model of depression (Willner, 2017). Mice in the no-CUS condition were housed in a separate colony room to avoid the transfer of olfactory cues from CUS-exposed mice (Brechbuhl et al., 2013), and left undisturbed with the exception of daily injections and weekly cage changing.

**Table 1.** Stressors used in the Chronic Unpredictable Stress paradigm.

| Stressor | Description |
| --- | --- |
| Wet bedding | 500 ml of tap water in home cage for 3 hours |
| No bedding | Bedding removed from home cage for 3 hours |
| Soiled cage | Placed in a cage soiled by other mice for 3 hours |
| Tail bleed | Tail blood collection via needle poke |
| Restraint | Placed in well ventilated 50ml falcon tube for 1 hour |
| Cage tilting | Cage tilted at a 45° angle for 4 hours |
| Cage rotation | Housing with unfamiliar mice for 3 hours |
| Social isolation | Overnight social isolation, 18 hours |
| Elevated platform | Mice placed individually on a clear plexiglass elevated platform for 5 minutes |
| Lights off | Lights off for 3 hours during the light cycle |
| Continuous lighting | Continuous room lighting for two consecutive night cycles |
| Stroboscopic light | Flashing light (1 flash/sec) in an otherwise dark colony room during the light cycle |
| Noise | White noise from radio for 1 hour |

### Behavioural Testing

All mice were assessed on a battery of tests starting the day after CUS. One test was applied per day, and testing occurred in the order depicted in **Fig. 1**. Mice were acclimatized to testing rooms for 1 hour prior to each test. All behavioural scoring was performed by experimenters blinded to group assignment.

#### Forced Swim Test

The forced swim test (FST) was used according to previous methods (Can et al., 2012) to assess stress-coping behaviour which is sensitive to estrogens and exposure to stressors (Monteiro et al., 2015; Rocha et al., 2005). Briefly, under dim light conditions, mice were individually subjected to a 6-minute swimming session in a 4L glass beaker containing clean water at 25 ± 0.5°C, filled to a depth of 15 cm. The test was filmed, and time spent in passive-coping behaviour across the test was scored. Mice were considered to display passive-coping behaviour when immobile, with the absence of any movements except those necessary to remain afloat or keep the head above water.

#### Tail Suspension Test

The tail suspension test (TST) was used as an additional test of stress-coping behaviour, and performed according to standard methods (Can et al., 2011). Briefly, mice were suspended by the tail using laboratory tape (Fisherbrand^TM^ labeling tape) placed 2-3 mm from the tip of the tail, in a 3-walled rectangular chamber. Prior to suspension, and to prevent tail-climbing behaviour which is common in this strain, tails were passed through a 4cm hollow rubber tubing to cover a portion of the tail, as per previous methods (Can et al., 2011). The 6-minute duration of the test was filmed, and percent time spent immobile was considered an index of passive-coping behaviour.

#### Sucrose preference test

Anhedonia-like behaviour was assessed in the sucrose preference test which was adapted from published methods (Pothion et al., 2004). Mice were acclimatized to two water bottles in the home cage for 24 hours and individually housed approximately 4 hours prior to testing. Testing occurred during the dark phase (19:00-07:00) in which mice were presented with two bottles, one containing 1% sucrose and the other tap water. The right-left position of sucrose was counterbalanced between mice in each group. Sucrose preference was calculated as the percentage of sucrose consumed over total liquid consumed.

#### Novelty Suppressed feeding

Anxiety-like behaviour was assessed using the novelty suppressed feeding test (NSF), adapted from previously published methods (Samuels and Hen, 2011). Behaviour in this test is sensitive to chronic stress exposure and ovarian status (Mahmoud et al., 2016; Stedenfeld et al., 2011), and chronic antidepressant treatment produces an anxiogenic effect in a neurogenesis-dependent manner in males (Santarelli et al., 2003). Briefly, mice were deprived of food for 24 hours, then individually placed in one corner of an open arena (50x50x20cm) containing a froot loop^©^ in the centre, and latency to start feeding was measured. Testing was performed under bright light conditions, and the arena was cleaned with 70% EtOH between mice. If a mouse did not feed within 10 minutes, it was removed from the arena and given a latency of 600 seconds in the analysis. Mice were returned to their home cage following testing, and food consumed within 5 minutes in the home cage was measured to account for potential appetite differences. Mice were re-grouped after the test. To avoid a neophobic reaction to the food, froot loops^©^ were introduced to the home cage for 3 days prior to the test.

### Dexamethasone suppression and HPA axis stress reactivity tests

Disrupted HPA axis negative feedback inhibition is an endocrine hallmark of MDD (Ising et al., 2007; Stetler and Miller, 2011), and estrogens regulate HPA negative feedback function (Goel et al., 2014). Therefore, half the animals from each group were subjected to the dexamethasone (DEX) suppression test to assess the integrity of the glucocorticoid-dependent negative-feedback function of the HPA axis. DEX (i.p.; 100 ug/kg) was administered 15 minutes prior to the regular treatment (E2, DPN, PPT, or oil). Mice were subjected to a 30-min restraint stressor starting 90 minutes after DEX administration, and blood was collected via tail vein nick at the end of restraint. To investigate HPA axis reactivity to an acute stressor, the other half of animals from each group received a vehicle injection (0.9% saline) in place of DEX, with the remainder of the procedure (restraint and blood collection) being identical. Blood samples were placed at 4°C overnight, then centrifuged for 15 min at 1000 x g and serum aliquots were stored at -20°C until processing.

### Tissue harvesting and processing

Mice were euthanized approximately 2 hours after the DEX/HPA axis stress reactivity test. To obtain fixed brain tissue for immunohistochemistry, approximately half the mice in each group were deeply anesthetized with sodium pentobarbital (i.p.) then perfused with 10mL of cold 0.9% saline followed by 20mL of cold 4% paraformaldehyde (PFA). Brains were extracted immediately and post-fixed at 4°C in 4% PFA for 24 hours, then transferred to a 30% sucrose solution (in 0.1 M Phosphate Buffer; pH 7.4) and kept at 4°C until sectioning. To obtain fresh tissue for electrochemiluminescence immunoassays, the other half of mice in each group were euthanized by rapid decapitation and brains were removed immediately and micro-dissected on a cold surface. Specifically, the entire rostral-caudal extent of the hippocampus was dissected, and the frontal cortex was collected anterior to the genu of the corpus callosum (Bregma 1.42mm). Tissue was flash-frozen on dry ice and stored at -80°C until homogenized. Blood was collected via cardiac puncture from perfused mice, and trunk blood was collected from decapitated mice. In both cases, blood was collected into EDTA-coated tubes, centrifuged for 15 minutes at 4°C at 1000 x g, then plasma aliquots were stored at -80°C until processing.

### Electrochemiluminescence immunoassays

In preparation for electrochemiluminescence immunoassays (ECLIAs), frontal cortices and hippocampi were homogenized individually in cold lysis buffer (200µl, 150µL, and 400µL, respectively) using an Omni Bead Ruptor (Omni international, Kennesaw, GA). Homogenates were immediately centrifuged at 2600rpm and 4°C for 10 minutes, and aliquots were stored at -80°C until processing. ECLIA kits from Meso Scale Discovery (MSD; Rockville MD) were used for the quantification of cytokines and PSD-95. All assays were performed in accordance with manufacturer’s instructions, with samples run in duplicates. Plates were read using a Sector Imager 2400 (MSD; Rockville MD), and data were analyzed using the Discovery Workbench 4.0 software (MSD). ECLIA values were normalized to total protein concentrations, which were measured in tissue homogenates in triplicates using the Pierce Micro BCA Protein Assay Kits (ThermoFisher Scientific) according to manufacturer instructions.

#### Cytokine quantification

Because inflammation is implicated in MDD (Dowlati et al., 2010; Enache et al., 2019; Haapakoski et al., 2015), several immune mediators were measured in the hippocampus and frontal cortex. The V-PLEX Proinflammatory Panel 1 kit (mouse) from MSD (catalogue no. K15048D) was used, which allows for the concurrent quantification of Interleukin-1beta (IL-1β), Interleukin-2 (IL-2), Interleukin-4 (IL-4), Interleukin-6 (IL-6), Interleukin-10 (IL-10), Interferon-gamma (IFN-γ), tumor necrosis factor-α (TNF-α), chemokine (C-X-C motif) ligand 1 (CXCL1), Granulocyte-macrophage colony-stimulating factor (GM-CSF), and vascular endothelial growth factor (VEGF), thus provides a comprehensive picture of the inflammatory state. The following lower limits of detection (LLODs) were observed: IL-1β: 0.74-0.88 pg/ml; IL-2: 2.1-2.6 pg/ml; IL-4: 0.35-0.45 pg/ml; IL-6: 5.6-6.2 pg/ml; IL-10: 1.1-1.8 pg/ml; IFN-γ: 0.07-0.11 pg/ml; TNF-α: 0.88-1.32 pg/ml; CXCL1: 0.17-0.36 pg/ml; GM-CSF: 0.09-0.13 pg/ml; and VEGF: 0.31-0.38 pg/ml. Values below the LLOD were assigned 0pg/mL, as published previously (Bodnar et al., 2017; Eid et al., 2019a).

#### PSD-95 quantification

PSD-95 was quantified because its expression is regulated by E2, DPN and PPT in the hippocampus of female rats and mice (Li et al., 2004; Waters et al., 2009) and because it’s reduced in the PFC of individuals with MDD (Feyissa et al., 2009) and with chronic stress exposure in rodents (Kallarackal et al., 2013; Kim and Leem, 2016; Pacheco et al., 2017). PSD-95 protein expression was quantified in hippocampus and frontal cortex samples using a PSD-95 kit from MSD (catalogue no. K150QND).

### Immunohistochemistry

For immunohistochemical staining, PFA-fixed brains were sliced into 30µm coronal sections using a Leica SM2000R Microtome (Richmond Hill, Ontario, Canada). Sections were stored at -20 °C in a cryoprotective solution containing 20% glycerol (Sigma-Aldrich, St. Louis, MO, USA) and 30% ethylene glycol (Sigma-Aldrich) in 0.1 M phosphate-buffer (PB, pH 7.4). Tissue was rinsed in 0.1M phosphate-buffered saline (PBS; pH 7.4; 3 x 10 min) to remove the cryoprotectant prior to staining and between all incubations. Unless otherwise noted, all incubations were performed on a rotator and at room temperature.

#### BrdU/NeuN

We examined the co-expression of BrdU with the neuronal marker NeuN to determine the proportion of BrdU immunoreactive (ir) cells that are neurons. Sections incubated for 24 hours at 4°C in a primary antibody solution containing mouse anti-NeuN (catalogue no. MAB377; Millipore, Billerica, MA, USA) diluted 1:250 in 0.3% Triton-X and 3% normal donkey serum (NDS) in 0.1M PBS. Sections were then transferred to a secondary antibody solution containing 1:200 donkey anti-mouse Alexa Fluor 488 (Invitrogen, Burlington, ON, Canada) and incubated for 24 hours at 4°C. Sections were fixed in 4% PFA (in 0.1 PBS; 1 x 10 mins) followed by 0.9% saline (2 x 10 mins), before being transferred to 2N HCL for 30 mins at 37°C. Sections were then incubated for 24 hours at 4°C in a primary antibody solution containing rat anti-BrdU (catalogue no. ab6326; Abcam, Cambridge, UK) diluted 1:500 in 0.3% Triton-X and 3% NDS in 0.1M PBS. Sections were transferred to a secondary antibody solution containing 1:500 donkey anti-rat Alexa Fluor 594 (Jackson ImmunoResearch) in 0.1M PBS for 24 hours. Sections were finally mounted onto glass slides (Superfrost Plus; Fisher scientific, Pittsburgh, PA, USA) and cover-slipped with polyvinyl alcohol (PVA)-DABCO (Sigma-Aldrich).

#### Ionized calcium binding adaptor molecule-1 (Iba-1)

Iba-1 is a calcium-binding protein commonly utilized as a microglial marker (Korzhevskii and Kirik, 2016). Microglia were investigated as positron emission tomography studies provide evidence of increased microglial activation in MDD, and this effect is associated with illness duration and correlated with depression severity (Setiawan et al., 2018, 2015). Hippocampal and ventromedial prefrontal cortex (vmPFC) sections were incubated for 25 min in 0.3% hydrogen peroxide (in dH_2_O) then blocked for 1 h with 10% normal goat serum (NGS) and 0.5% Triton-X in 0.1M PBS. Sections were transferred to a primary antibody solution containing rabbit anti-Iba-1 (1:1000; catalogue no. 019-19741; Wako, Osaka, Japan) in 10% NGS and 0.4% Triton-X in 0.1M PBS and incubated at 4°C for 20 hours. Next, sections were incubated in a secondary antibody solution for 1 hour, consisting of 1:500 biotinylated goat anti-rabbit (Vector Laboratories) in 2.5% NGS and 0.4% Triton X in PBS. Sections were then incubated for 1 h in ABC (Elite kit; 1:50, Vector Laboratories) in 0.4% Triton-X in PBS, then immunoreactants were visualized with a 7 min diaminobenzidine (DAB) reaction (Vector Laboratories). Sections were mounted on glass slides, allowed to dry, then dehydrated in increasing graded ethanol solutions, cleared with xylene, and cover-slipped with Permount (Fisher Scientific).

### Microscopy, cell quantification, and optical density

BrdU- immunoreactive (ir) cell quantification was performed by an experimenter blinded to group assignment under a 100x objective on an Olympus BX51 brightfield microscope equipped with epifluorescence. A representative photomicrograph is depicted in **Fig. 4C**. BrdU-ir cells were exhaustively quantified in every 10^th^ section of the hippocampus specifically within the granule cell layer (GCL) and the subgranular zone (SGZ), defined as the 50µm band between the GCL and hilus. Cells were quantified separately in the dorsal and ventral hippocampus as these regions have distinctive functions (Fanselow and Dong, 2010) Raw counts were multiplied by 10 to obtain an estimate of the total number of BrdU-ir cells in the hippocampus (Mahmoud et al., 2016a; Pan et al., 2012). BrdU-ir cells were examined for NeuN co-expression to estimate the proportion of BrdU-ir cells that are neurons (calculated as BrdU/NeuN co-labeled cells divided by the total number of BrdU-ir cells). Then, to obtain a neurogenesis index for each animal, the estimated total number of BrdU-ir cells was multiplied by the ratio of BrdU/NeuN co-labeled cells.

Optical density of Iba-1 expression was measured using ImageJ in images obtained using CellSens software (Olympus) at 10x with fixed gain and exposure settings, as we have done previously (Wainwright et al., 2016; Workman et al., 2015). Six sections were sampled per animal, including two sections each for the dorsal hippocampus (Bregma -1.66 to -2.35mm), ventral hippocampus (Bregma - 3.18 to -3.58mm), and vmPFC (Bregma 1.54 to 1.94mm, with infralimbic (IL) and prelimbic (PL) regions sampled separately). Background grey value was obtained in each image from an average of open ellipses placed randomly in 10 non-immunolabeled areas, and threshold for immunoreactivity was set at 3-times the mean background grey value. In hippocampal slices, the GCL, SGZ and an approximately 50μm band of the molecular layer (ML) were traced and mean grey value per area was calculated. In vmPFC slices, mean grey value per area was measured in two 0.85mm^2^ regions placed within the IL and PL regions.

### Vaginal cytology for estrous cycle staging

Behaviour, immune, and neural measures can be affected by estrous cycle stage (Beagley and Gockel, 2003; Meziane et al., 2007; Woolley et al., 1990). Therefore, vaginal cells were collected by lavage from sham-operated mice on all behavioural testing days and prior to euthanasia, and estrous cycle stage was determined according to previous methods (Cora et al., 2015). To account for potential effects of the procedure, all ovariectomized mice were also lavaged.

### Radioimmunoassays for corticosterone quantification

Corticosterone concentrations were measured in serum samples obtained from HPA axis stress reactivity and DEX suppression tests. Corticosterone double-antibody radioimmunoassay kits (MP biomedicals, Solon, OH) were used according to manufacturer’s instructions and the Inter- and intra-assay coefficients of variation were < 10%.

### Statistical analyses

Statistica software (Tulsa, OK) was used for all analyses. Behavioural measures (FST, TST, NSF, and SPT), cytokines, and CORT concentrations were each analyzed by factorial analysis of variance (ANOVA) with treatment (Sham, OVX, E2, DPN, PPT) and stress condition (CUS, no-CUS) as the between-subject factors. Neurogenesis index, proportion of of BrdU/NeuN co-labeled cells, PSD-95 expression, and Iba-1 optical density were each analyzed using repeated measures ANOVA with treatment (Sham, OVX, E2, DPN, PPT) and stress condition (CUS, no-CUS) as the between-subject factors and brain region (dorsal and ventral hippocampus, or hippocampus and frontal cortex) as the within-subject factors. Given that 10 cytokines were measured in each brain region, Principal Component Analyses (PCAs) were used as a dimensionality reduction tool (Jolliffe, 2002; Ringner, 2008). PCA reduces the number of potentially correlated variables (in this case cytokines) into a small set of uncorrelated variables called Principal Components (PCs). These components are ordered such that the first PC (PC1) accounts for most of the variance within the data. This approach was used to derive information about the amount of variance in the data that can be accounted for by potential cytokine networks. Because PCA does not take into account group membership, ANOVAs were subsequently applied to individual PC scores as we have published previously (Eid et al., 2019a), in order to explore the effects of treatments and CUS exposure on cytokine networks. DEX was used as a covariate in all cytokine analyses as DEX was administered to half the animals on tissue collection day, and because glucocorticoids have anti-inflammatory effects (Bereshchenko et al., 2018). Body mass was used as a covariate when analyzing immobility in FST. Estrous cycle stage was used as a covariate for all analyses. Covariate effects are only reported when significant. Neuman-Keul’s comparisons were used for post-hoc analyses and any *a priori* comparisons employed were subjected to Bonferroni correction. Outliers that fell more than 2.5 standard deviations away from the mean were removed from analyses. Significance was set at α=0.05. Were appropriate, effect sizes are given as partial η^2^ or Cohen’s d.

## Results

### TST: 1) CUS increased immobility, and this effect was primarily driven by ER agonists groups. 2) Ovariectomy increased immobility, and this effect was prevented by estrogenic treatments only under no-CUS Conditions

As expected, CUS significantly increased immobility in the TST (main effect of CUS condition: F(1, 111)=11.96, p<0.001, η^2^_p_=0.097; Fig. 2A1). This effect of CUS to increase immobility was clearly driven by the ER agonist groups (PPT and DPN; both p’s < 0.0052, Cohen’s *d* = 0.88-1.1, relative to no-CUS counterparts; **Fig. 2A2-3**), and to a lesser extent by the E2 group (p=0.10, relative to no-CUS counterparts; a priori comparisons: treatment by CUS condition interaction: p=0.19). Further, ovariectomy increased immobility in the TST relative to Sham and DPN groups (p’s <0.03, Cohen’s *d* = 0.7-1.0; main effect of treatment: F(4, 111)=3.06, p=0.02, η^2^_p_ = 0.099; **Fig. 2A2**). Upon closer examination, this ovariectomy-induced increase in immobility was prevented by E2, DPN, and PPT treatments under no-CUS conditions only (p’s <0.04; *a priori* comparisons, treatment by CUS condition interaction: p=0.19; **Fig. 2A3**). In an additional exploratory analysis of TST data, we performed a median split and categorized mice into low immobility or “resilient”, versus high immobility or “susceptible” phenotypes, adapted from previous reports (Wells et al., 2017). We then performed a Chi-square test to compare the frequency of mice that were susceptible versus resilient between groups under CUS conditions: χ^2^(9) = 16.17, p = 0.06 (frequencies reported in **Table 2**). In the CUS exposed mice only, OVX mice displayed a bias towards a susceptible phenotype, whereas sham-operated mice displayed a bias towards a resilient phenotype. Further, a bias towards a susceptible phenotype is seen in PPT-treated groups, whereas E2- and DPN-treated groups showed an even split of resilient and susceptible mice after CUS.

**Table 2.** Percent time spent immobile in the forced swim test, percent sucrose preference (± standard error of the mean), and frequency of resilient versus susceptible in the tail suspension test.

| Group | % time spent immobile in<br>FST | % sucrose preference | # of resilient in TST<br>(low immobility) | # of susceptible in TST<br>(high immobility) |
| --- | --- | --- | --- | --- |
| **no-CUS** |  |  |  |  |
| Sham | 47.75 ± 3.49 | 82.24 ± 6.49 | 7 | 3 |
| OVX | 45.69 ± 5.11 | 84.62 ± 3.37 | 4 | 10 |
| E2 | 51.60 ± 4.97 | 89.64 ± 1.51 | 9 | 5 |
| DPN | 48.27 ± 4.28 | 87.79 ± 2.25 | 8 | 3 |
| PPT | 41.35 ± 3.81 | 86.26 ± 1.93 | 9 | 5 |
| **CUS** |  |  |  |  |
| Sham | 41.92 ± 3.49 | 90.92 ± 2.26 | 7 | 3 |
| OVX | 42.38 ± 3.86 | 90.98 ± 0.81 | 4 | 9 |
| E2 | 47.84 ± 2.97 | 89.86 ± 3.84 | 6 | 7 |
| DPN | 41.72 ± 3.35 | 92.75 ± 1.30 | 6 | 7 |
| PPT | 42.94 ± 4.15 | 81.06 ± 3.95 | 3 | 10 |

No significant group differences were observed. CUS, chronic unpredictable stress; Sham, sham-operated; OVX, ovariectomized; E2, estradiol; DPN, diarylpropionitrile; PPT, propylpyrazole-triol; FST, forced swim test; NSF, novelty suppressed feeding, TST; tail suspension test.

### Ovariectomy reduced latency to feed in NSF regardless of CUS exposure, and this effect was prevented by E2 treatment

Regardless of CUS condition, OVX significantly reduced latency to feed in NSF in comparison to Sham (p=0.029, Cohen’s *d* = 0.63; main effect of treatment: F(4, 111)=3.99, p=0.005, η^2^_p_ = 0.13; **Fig 2B**). This effect of OVX to reduce latency to feed was prevented by E2 (p=0.92, relative to Sham) but not by DPN or PPT treatments (p’s <0.05, relative to Sham). There was no significant main effect of CUS condition nor a treatment by CUS condition interaction (p’s >0.15).

**Figure 2.**
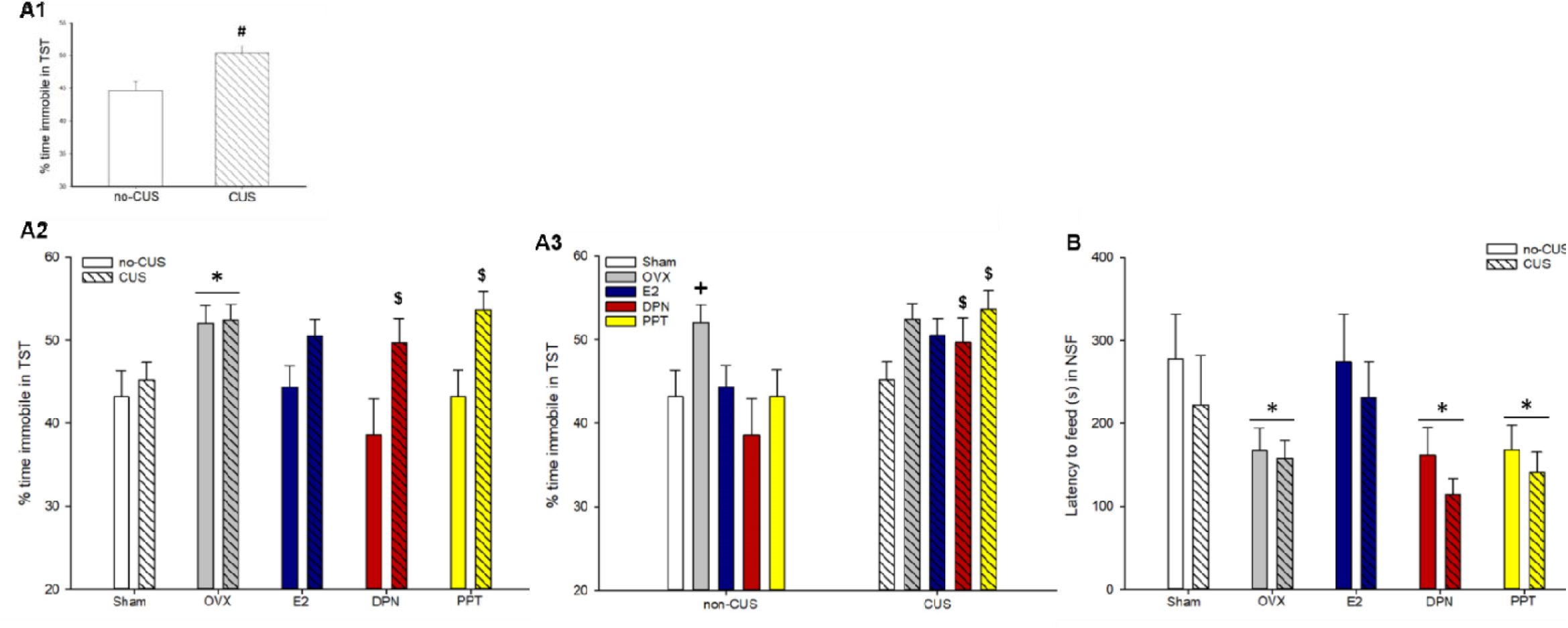
Percentage of time spent immobile in the tail suspension test (A1-3), and latency to feed in the novelty suppressed feeding test (B). Figures **A2** and **A3** depict the same data, organized differently across the x-axis for clarity of effects. **(A1)** CUS significantly increased immobility in TST; # indicates p<0.001, main effect of CUS condition. **(A2, A3)** The effect of CUS to increase immobility was primarily driven by DPN- and PPT-treated groups; $ indicate p’s <0.0052, significantly higher than no-CUS treatment counterparts. **(A2)** Ovariectomy significantly increased immobility, * indicates p <0.03, relative to Sham and DPN. **(A3)** the effect of ovariectomy to increase immobility was prevented by all estrogenic treatments only under non-CUS conditions only; **+** indicates p’s <0.05, relative to all non-CUS groups. **(B)** Ovariectomy significantly decreased latency to feed in NSF, and this effect was prevented by E2 treatment only, * indicate p <0.05, relative to sham. TST, tail suspension test; NSF, novelty suppressed feeding; CUS, chronic unpredictable stress; OVX, ovariectomized; E2, estradiol; DPN, diarylpropionitrile; PPT, propylpyrazole-triol. Data in means + standard error of the mean.

### There were no significant group differences in FST immobility or sucrose preference

There were no significant group differences in % time spent immobile in the FST or % sucrose preference (p’s >0.11 for main effects of CUS condition and treatment and their interaction; **Table 2**).

### E2 treatment reduced CORT response to acute restraint stress

In the HPA axis stress reactivity test, E2 treatment reduced post-restraint stress CORT concentrations relative to all other treatment groups (p’s <0.025, Cohen’s *d* = 0.6-1.8; main effect of treatment: F(4, 49)=3.96, p=0.007, η^2^_p_ = 0.24; **Fig. 3A**). There was no significant main effect of CUS condition nor a treatment by CUS condition interaction (p’s >0.4).

**Figure 3.**
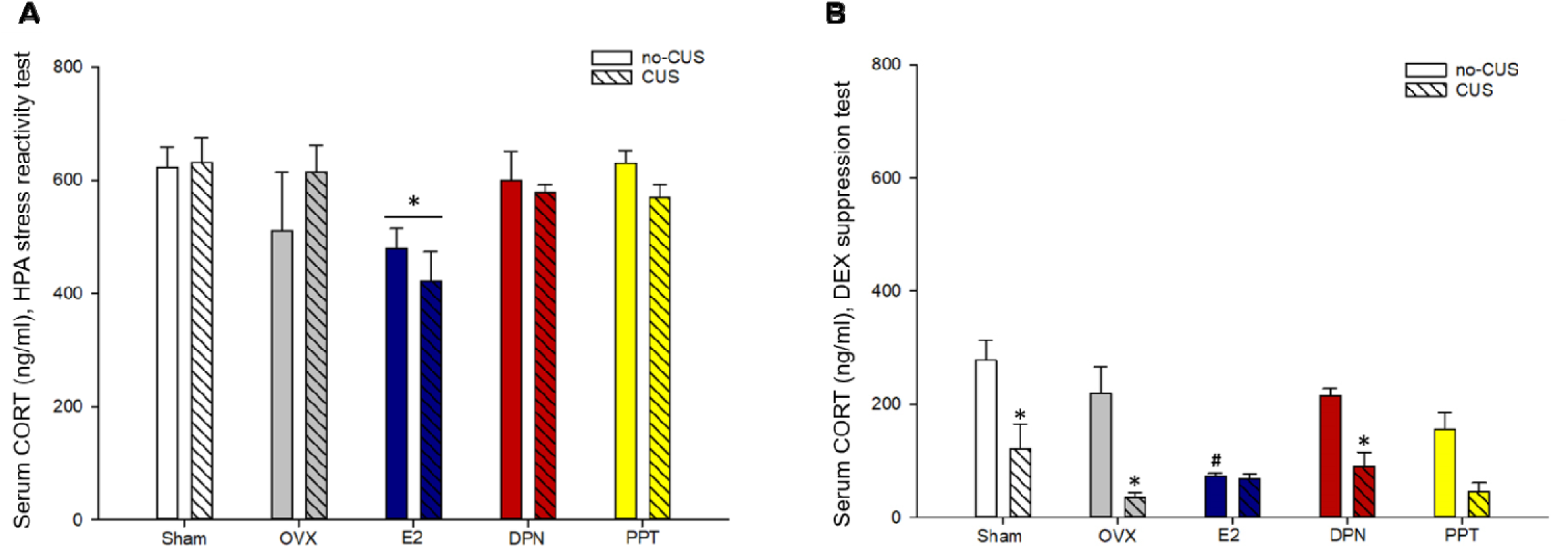
Serum corticosterone concentrations in the hypothalamic-pituitary-adrenal (HPA) axis stress reactivity test (A) and dexamethasone (DEX) suppression test (B). **(A)** E2 treatment reduced serum corticosterone concentrations immediately following an acute restraint stressor; * indicates p<0.025 relative to all other treatment groups. **(B)** CUS exposure significantly enhanced dexamethasone suppression of corticosterone in Sham, OVX and DPN groups, and a similar but non-significant trend was detected with PPT treatment (* indicates p<0.02 relative to no-CUS counterparts). Further, under no-CUS conditions E2 treatment reduced corticosterone concentrations (**#** indicates p <0.01 relative to Sham, OVX, and DPN treated no-CUS groups). CUS, chronic unpredictable stress; OVX, ovariectomized; E2, estradiol; DPN, diarylpropionitrile; PPT, propylpyrazole-triol. Data in means + standard error of the mean.

### CUS enhanced the dexamethasone suppression of CORT in most treatment groups

In the DEX suppression test, CUS exposure significantly reduced post-restraint stress CORT concentrations in Sham (p=0.0015, Cohen’s *d* = 1.8), OVX (p=0.00057, Cohen’s *d* = 2.0), and DPN (p=0.011, Cohen’s *d* = 2.5) groups, with a trend in PPT (p=0.06), but not in E2 (p=0.9; **Fig 3B**; treatment by CUS condition interaction: F(4, 51)=3.42, p=0.015, η^2^_p_ = 0.21). Further, E2 treated mice had sigificantly lower CORT concentrations relative to all treatment groups in the no-CUS condition (p<0.01; **Fig 3B**) except PPT (p=0.15). There were also significant main effects of treatment and stress condition (p’s <0.001).

### CUS exposure reduced the proportion of BrdU-ir cells that expressed NeuN in ventral dentate gyrus

In the ventral dentate gyrus only, CUS exposure reduced the proportion of BrdU/NeuN co-labeled cells (p = 0.04, Cohen’s *d* = 0.4; region by CUS condition interaction: F(1, 53)=5.73, p=0.02; **Table 3**).

**Table 3.**
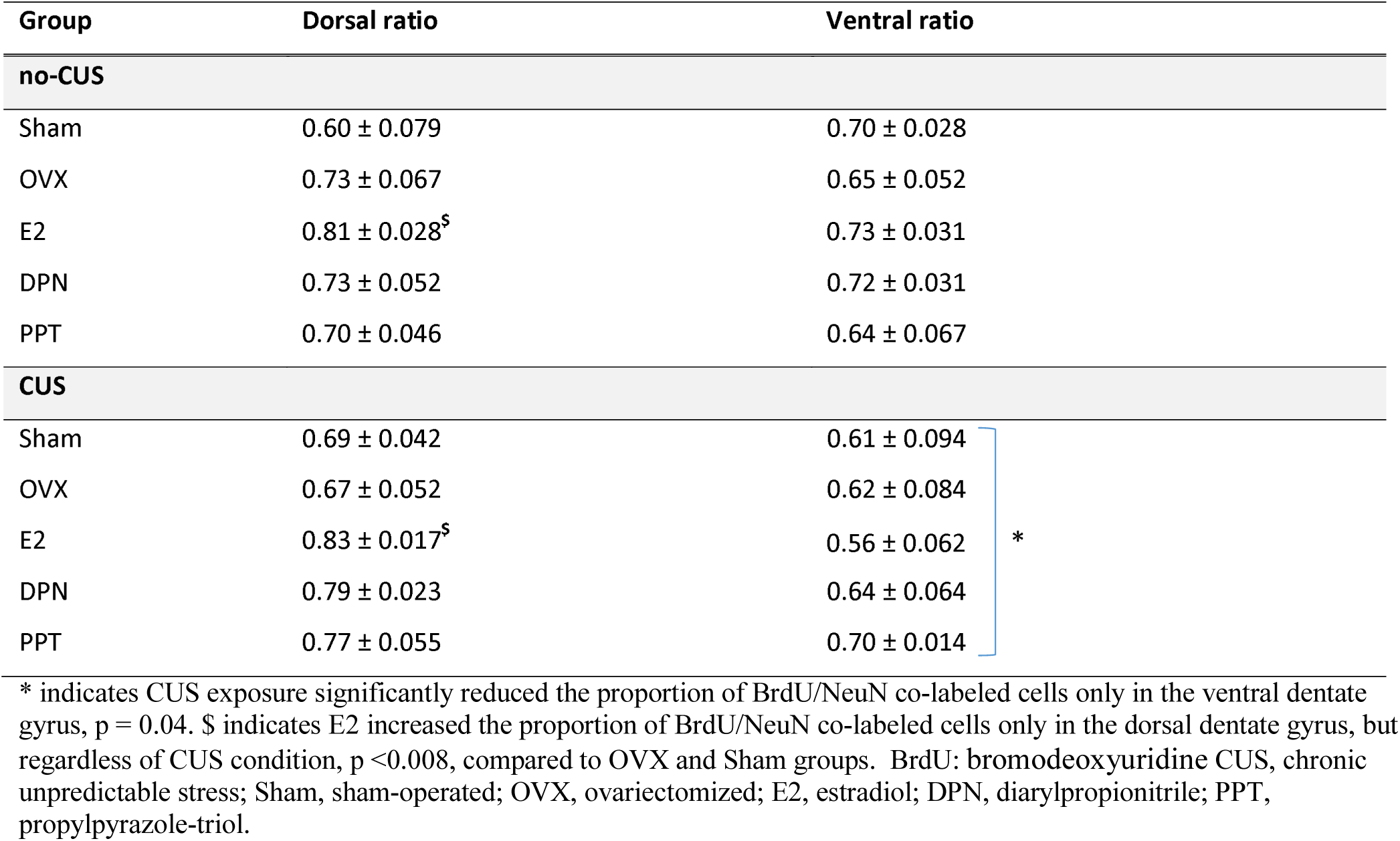
Proportion of BrdU/NeuN co-labeled cells in dorsal and ventral dentate gyrus (mean ± standard error of the mean)

| Group | Dorsal ratio | Ventral ratio |
| --- | --- | --- |
| **no-CUS** |  |  |
| Sham | 0.60 ± 0.079 | 0.70 ± 0.028 |
| OVX | 0.73 ± 0.067 | 0.65 ± 0.052 |
| E2 | 0.81 ± 0.028<sup>\$</sup> | 0.73 ± 0.031 |
| DPN | 0.73 ± 0.052 | 0.72 ± 0.031 |
| PPT | 0.70 ± 0.046 | 0.64 ± 0.067 |
| **CUS** |  |  |
| Sham | 0.69 ± 0.042 | 0.61 ± 0.094 |
| OVX | 0.67 ± 0.052 | 0.62 ± 0.084 |
| E2 | 0.83 ± 0.017<sup>\$</sup> | 0.56 ± 0.062 |
| DPN | 0.79 ± 0.023 | 0.64 ± 0.064 |
| PPT | 0.77 ± 0.055 | 0.70 ± 0.014 |

\* indicates CUS exposure significantly reduced the proportion of BrdU/NeuN co-labeled cells only in the ventral dentate gyrus, $p = 0.04$ . \$ indicates E2 increased the proportion of BrdU/NeuN co-labeled cells only in the dorsal dentate gyrus, but regardless of CUS condition, $p < 0.008$ , compared to OVX and Sham groups. BrdU: bromodeoxyuridine CUS, chronic unpredictable stress; Sham, sham-operated; OVX, ovariectomized; E2, estradiol; DPN, diarylpropionitrile; PPT, propylpyrazole-triol.

Further, the proportion of BrdU/NeuN co-labeled cells was overall higher in the dorsal, compared to the ventral, dentate gyrus (main effect of region: F(1, 53)=14.508, p=0.0004. η^2^ =0.2). There was also a trend for a region by treatment interaction (p=0.08), where, intriguingly, E2 increased the proportion of dorsal BrdU-ir cells that also expressed NeuN, suggesting a promotion of cell fate to neuronal phenotype with chronic E2, regardless of stress condition (compared to OVX and Sham p’s <0.008; a priori comparisons). There were no other significant main effects or interactions (p’s >0.2).

### CUS reduced neurogenesis in the ventral dentate gyrus and E2 increased neurogenesis in the dorsal dentate gyrus

Irrespective of CUS exposure, E2 significantly increased neurogenesis index in the dorsal GCL only, relative to all other treatments (all p’s <0.0003, Cohen’s *d* = 0.7-1.2; region by treatment interaction: F(4, 53)=3.37, p=0.016, η^2^_p_ =0.2; **Fig 4A**). CUS exposure reduced neurogenesis in the ventral dentate gyrus (p=0.05, Cohen’s *d* = 0.5) but not in the dorsal dentate gyrus (p=0.28; region by CUS condition interaction: F(1, 53)=7.48, p=0.008, η^2^_p_ =0.12; **Fig 4B**). There were also significant main effects of treatment and region (p’s <0.04) but no other main effects or interactions (p’s >0.5).

**Figure 4.**
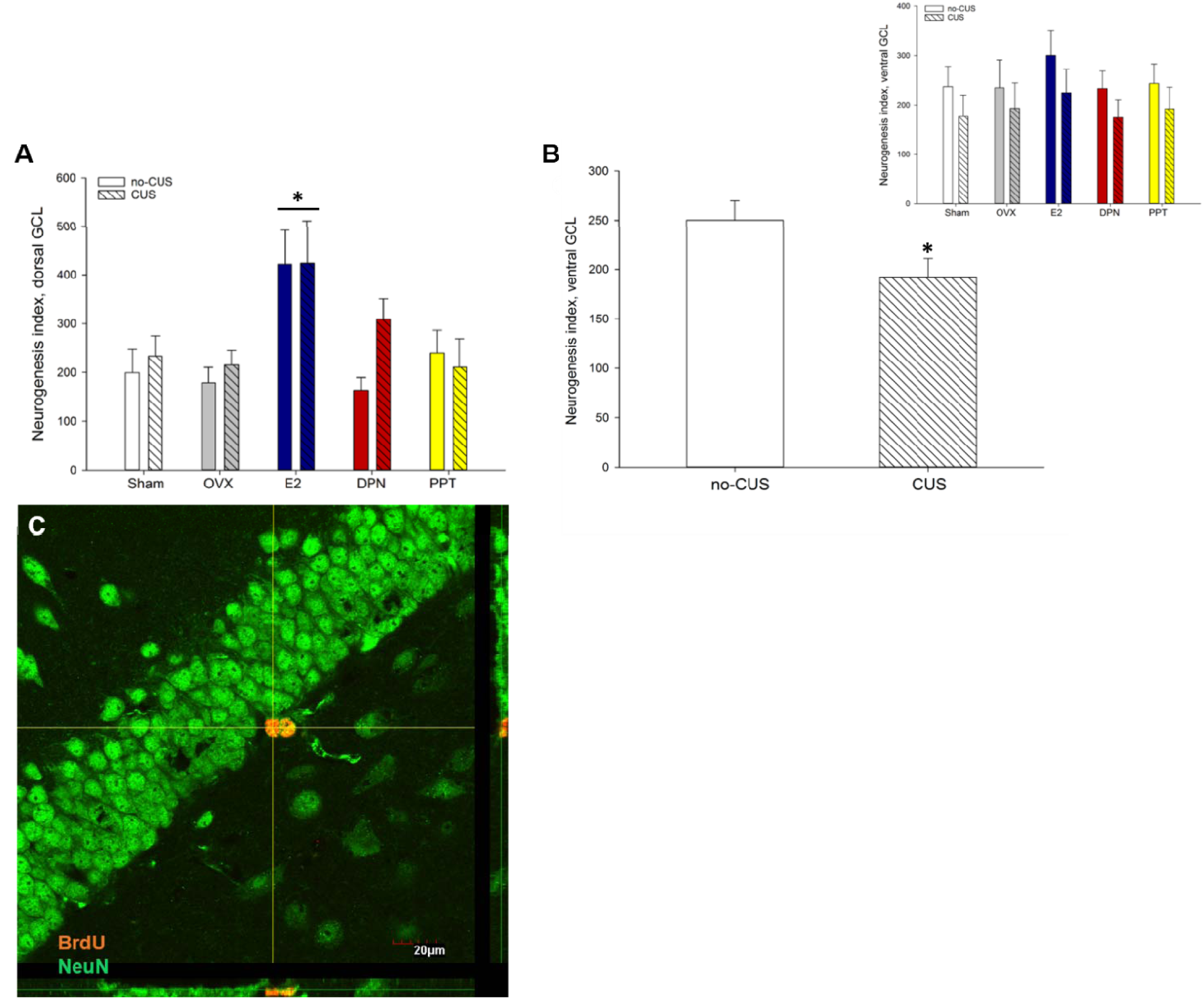
Neurogenesis index in the dorsal (A) and ventral (B) dentate gyrus. **(A)** Estradiol treatment significantly increased neurogenesis in the dorsal granule cell layer (GCL); * indicates p<0.0003 relative to all other treatment groups. **(B)** CUS exposure significantly reduced neurogenesis in the ventral GCL; **\*** indicates p = 0.05 relative to no-CUS. Inset graph depicts the data by treatment and CUS condition. **(C)** Representative photomicrographs of the granule cell layer (GCL) of the dentate gyrus, obtained at 60x magnification showing two BrdU-ir cells (orange) co-expressing NeuN (green). GCL, granule cell layer BrdU, bromodeoxyuridine; CUS, chronic unpredictable stress; OVX, ovariectomized; E2, estradiol; DPN, diarylpropionitrile; PPT, propylpyrazole-triol. Data in means + standard error of the mean.

### CUS exposure decreased PSD-95 expression in the hippocampus and frontal cortex, and this effect was observed primarily in DPN- and PPT-treated groups

CUS exposure significantly decreased PSD-95 expression in the hippocampus and frontal cortex (main effect of CUS condition: F(1, 45)=7.89, p=0.007, η^2^_p_ =0.15; **Fig 5A**). The effects of CUS to decrease PSD-95 were largely driven by PPT and DPN groups (p’s<0.05, relative to non-CUS counterparts; CUS condition by treatment interaction: F(4, 45)=2.67, p=0.044, η^2^_p_ =0.19). Further, PSD-95 expression was overall significantly higher in the hippocampus than the frontal cortex (main effect of region: F(1, 45)=98.18, p<0.001 η^2^_p_=0.69; **Fig 5B**). There were no other main effects or interactions (all p’s >0.1).

**Figure 5.**
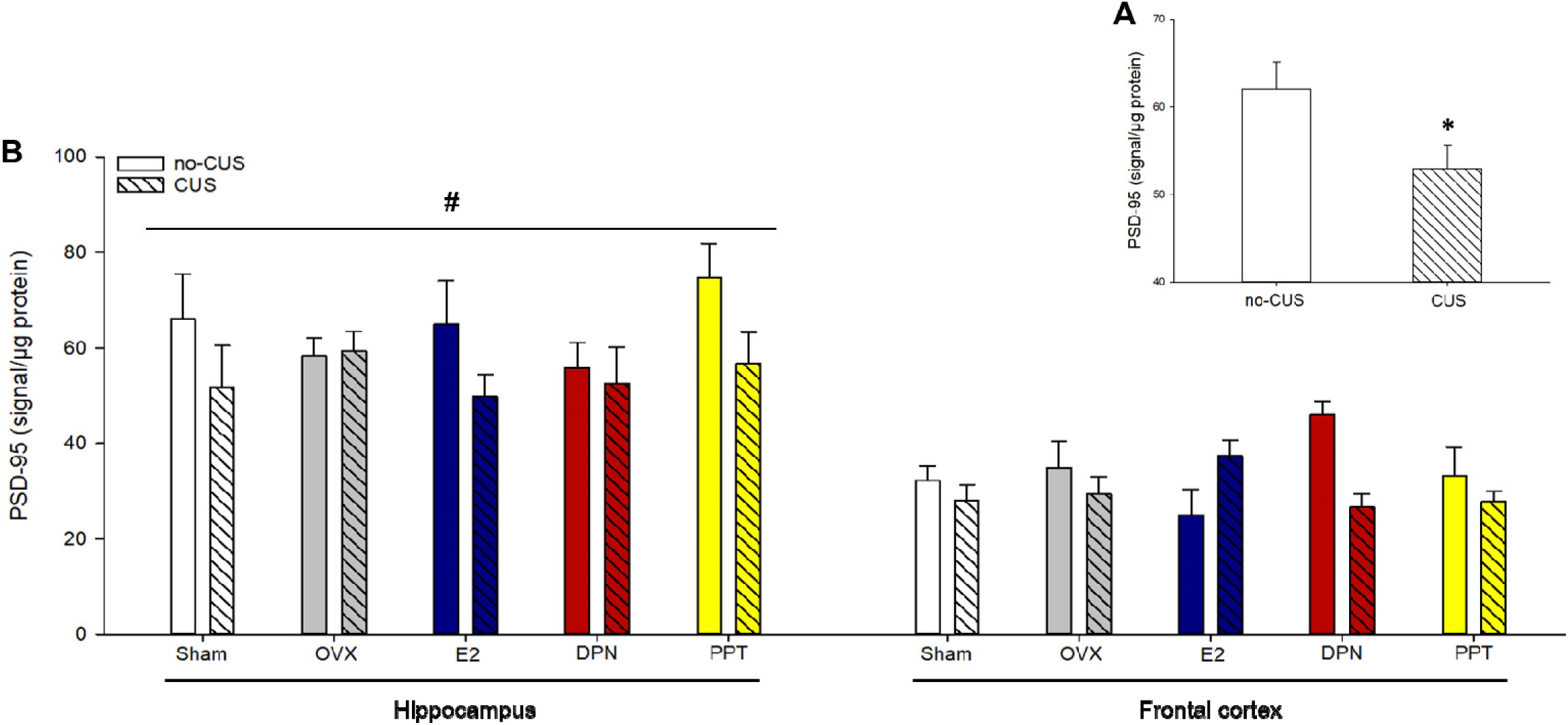
Postsynaptic density protein 95 (PSD-95) expression in the hippocampus and frontal cortex, normalized by total protein concentrations. **(A)** CUS exposure decreased PSD-95 expression in both regions; * indicate P=0.007, main effect of CUS condition. **(B)** PSD-95 expression was significantly higher in the hippocampus in comparison with the frontal cortex. # indicates p <0.001, main effect of region. PSD-95, postsynaptic density-95; CUS, chronic unpredictable stress; OVX, ovariectomized; E2, estradiol; DPN, diarylpropionitrile; PPT, propylpyrazole-triol. Data in means + standard error of the mean.

## Cytokine PCA results

### CUS increased PC1 scores in the hippocampus, an effect driven primarily by Sham and DPN-treated groups, and Sham-operated mice had lower PC2 scores

The model generated two principal components, accounting for 57.5% of the variance within the dataset, with variance explained by principal component 1 (PC1) = 39.7%, and PC2 = 17.8%. Factor loadings are shown in **Table 4**. ANOVA results reveal that CUS exposure increased PC1 scores (main effects of CUS condition: F(1, 53)=6.2920, p=0.015, η^2^_p_ =0.11; **Fig. 6A**), a significant covariate effect of DEX to reduce scores (p=0.002), but no significant main effect of treatment (p=0.45). The effect of CUS exposure to increase PC1 scores appears to be driven by Sham and DPN groups (p=0.027 and 0.017, relative to no-CUS counterparts; *a priori* comparisons, CUS condition by treatment interaction: F(4, 53)=1.63, p=0.18). PC2 scores were lower in Sham relative to all other treatment groups (p’s <0.02; main effect of treatment: F(4, 52)=3.39, p=0.016, η^2^_p_ =0.21; **Fig. 6B**), but this missed significance in comparison to PPT treatment (p=0.08). There was also trend toward significance for CUS to reduce PC2 scores (p=0.07), but no significant CUS condition by treatment interaction (F(4, 52)=1.3093, p=0.28).

**Figure 6.**
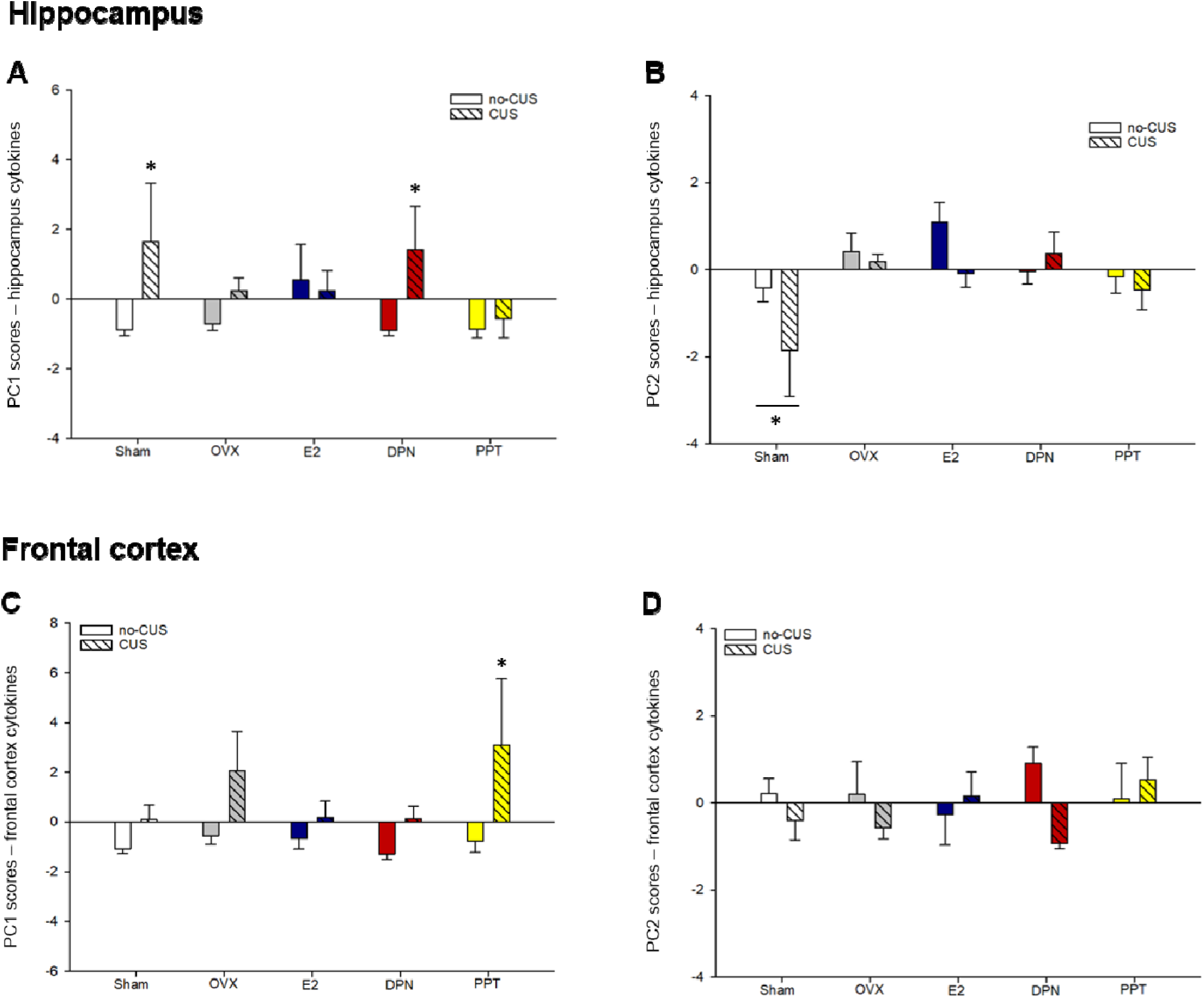
Principal component scores in hippocampus (A-B) and frontal cortex (C-D) analyses of cytokine data. **(A)** In the hippocampus, CUS exposure significantly increased principal component (PC) 1 scores, and this effect was driven by=Sham and DPN groups only; * indicate p<0.03, relative to no-CUS counterparts. **(B)** Sham-operated groups had significantly lower PC2 scores in the hippocampus; * indicates p’s <0.02, in comparison with all treatments except PPT (p=0.08). **(C)** In the frontal cortex, CUS exposure significantly increased PC1 scores, and this effect was driven by PPT treatment; * indicates p=0.003 relative to no-CUS counterparts. **(D)** There were no significant group differences in frontal cortex PC2 scores. PC, principal component; CUS, chronic unpredictable stress; OVX, ovariectomized; E2, estradiol; DPN, diarylpropionitrile; PPT, propylpyrazole-triol. Data in means + standard error of the mean.

**Table 4.** Principle component analyses loading table.

|  | Hippocampus |  | Frontal cortex |  |
| --- | --- | --- | --- | --- |
| Variable | PC1 | PC2 | PC1 | PC2 |
| GM-CSF | **0.52** | -0.18 | 0.20 | **0.81** |
| IFN-γ | **0.71** | 0.38 | **0.78** | 0.20 |
| IL-10 | **0.90** | -0.06 | **0.89** | 0.15 |
| IL-1 $\beta$ | **0.87** | -0.30 | **0.90** | -0.07 |
| IL-2 | **0.59** | **-0.63** | **0.80** | 0.14 |
| IL-4 | **0.64** | 0.30 | **0.73** | 0.15 |
| IL-6 | **0.41** | **0.56** | **0.91** | -0.05 |
| CXCL1 | 0.11 | **0.64** | -0.22 | **0.68** |
| TNF-α | **0.82** | -0.12 | **0.94** | -0.13 |
| VEGF | 0.15 | **0.52** | -0.37 | **0.76** |

### CUS increased PC1 scores in the frontal cortex, an effect driven primarily by PPT treatment

The model generated two principal components, accounting for 71.4% of the variance within the dataset, with variance explained by PC1 = 53.2% and PC2 = 18.2%. Factor loadings are shown in **Table 4**. ANOVA results show that CUS exposure significantly increased PC1 scores (main effect of CUS condition: F(1, 45)=13.43, p=0.00065, η^2^_p_ =0.23; **Fig. 6C**). This effect appears to be driven by PPT-treated mice (p=0.003; relative to no-CUS counterparts) with a trend toward significance in OVX (p=0.068; *a priori* comparisons; treatment by CUS condition interaction: F(4, 45)=1.12, p=0.36). There was also a significant covariate effect of DEX (p=0.024), but no significant main effect of treatment (p=0.14). PC2 scores were not significantly affected by treatment, CUS condition, nor their interaction (p’s >0.1; **Fig. 6D**), but there was a significant covariate effect of DEX (p=0.009).

### Individual cytokine results: CUS increased several cytokines in the hippocampus and frontal cortex, but the effects of estrogens were limited to VGEF and CXCL1 in the hippocampus

In the hippocampus, CUS exposure significantly increased TNF-α, IL-1β, and IL-10 concentrations (main effects of CUS: F(1, 53)=6.84, p=0.012, η^2^_p_=0.11; F(1, 51)=12.87, p=0.0007, η^2^_p_= 0.20; and F(1, 53)=7.38, p=0.0009, η^2^ =0.12, respectively; **Fig. 7A-C**), but decreased CXCL1 concentrations (F(1, 52)=4.01, p=0.05; **Fig. 7D**). However, IL-2, IL-4, IL-6, IFN-γ, VGEF, and GM-CSF were not significantly affected by CUS exposure (p’s>0.16). Estrogens only significantly influenced two proteins: VEGF and CXCL1. Specifically, the ERβ agonist, DPN, increased hippocampal VEGF concentration relative to Sham and PPT groups (p’s <0.03; main effect of treatment: F(4, 53)=3.55, p=0.012, η^2^=0.21; **Fig. 7E**), and E2 significantly increased hippocampal CXCL1 concentrations relative to Sham mice (p = 0.0045; main effect of treatment: F(4, 52)=3.57, p=0.012, η^2^ =0.22; **Fig. 7D**). There were no other significant main or interaction effects for any other cytokine in the hippocampus (p’s >0.083; **Table 5**). Finally, there was a significant covariate effect of DEX to reduce IL-1β, IL-6, IL-10, and TNF-α concentrations (p’s<0.05) and trends toward significance to reduce IL-2 (p=0.051) and GM-CSF (p=0.07) concentrations. DEX was not a significant covariate for concentrations of IFN-γ, VGEF, CXCL1, IL-4 (p’s >0.14).

**Figure 7.**
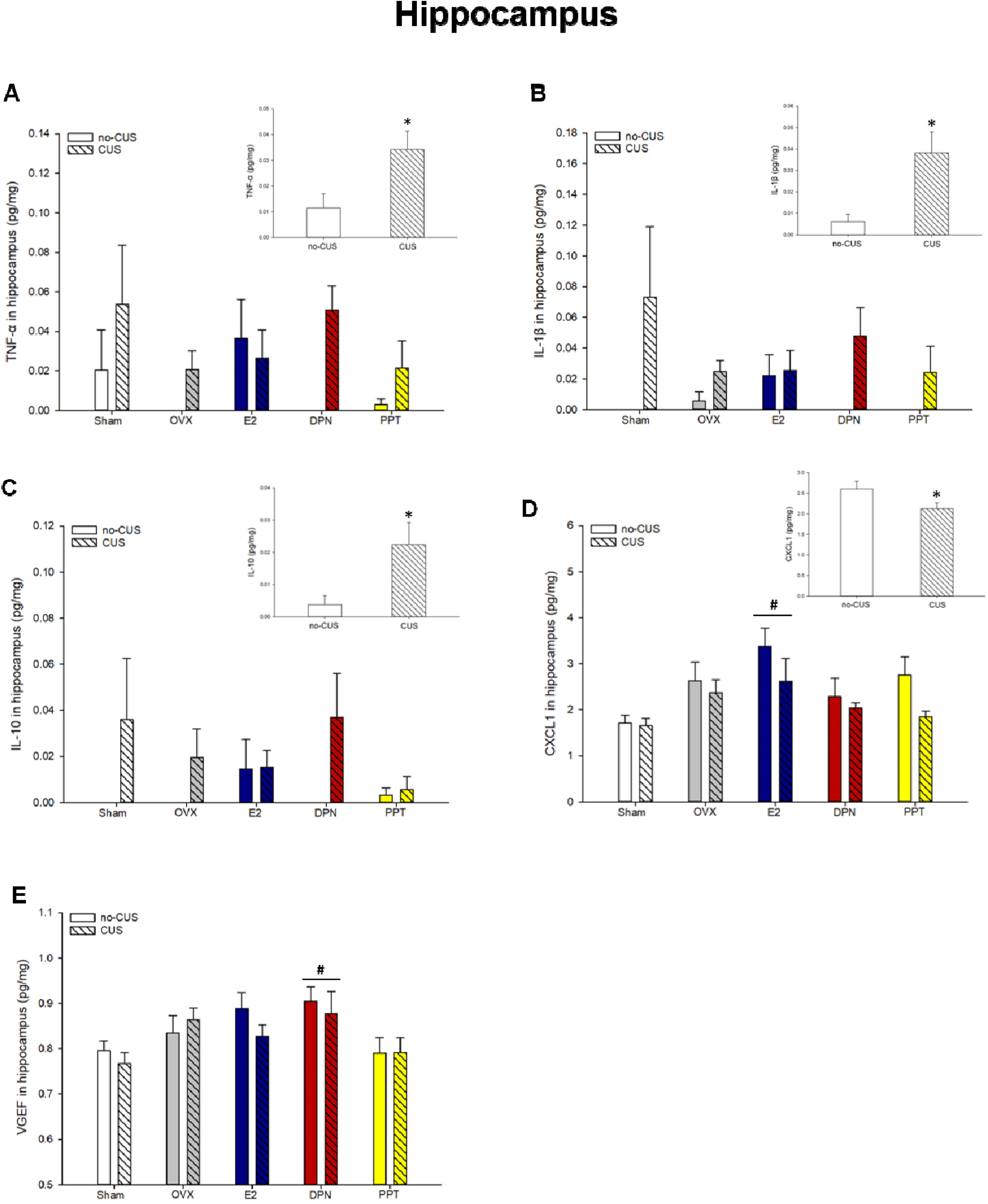
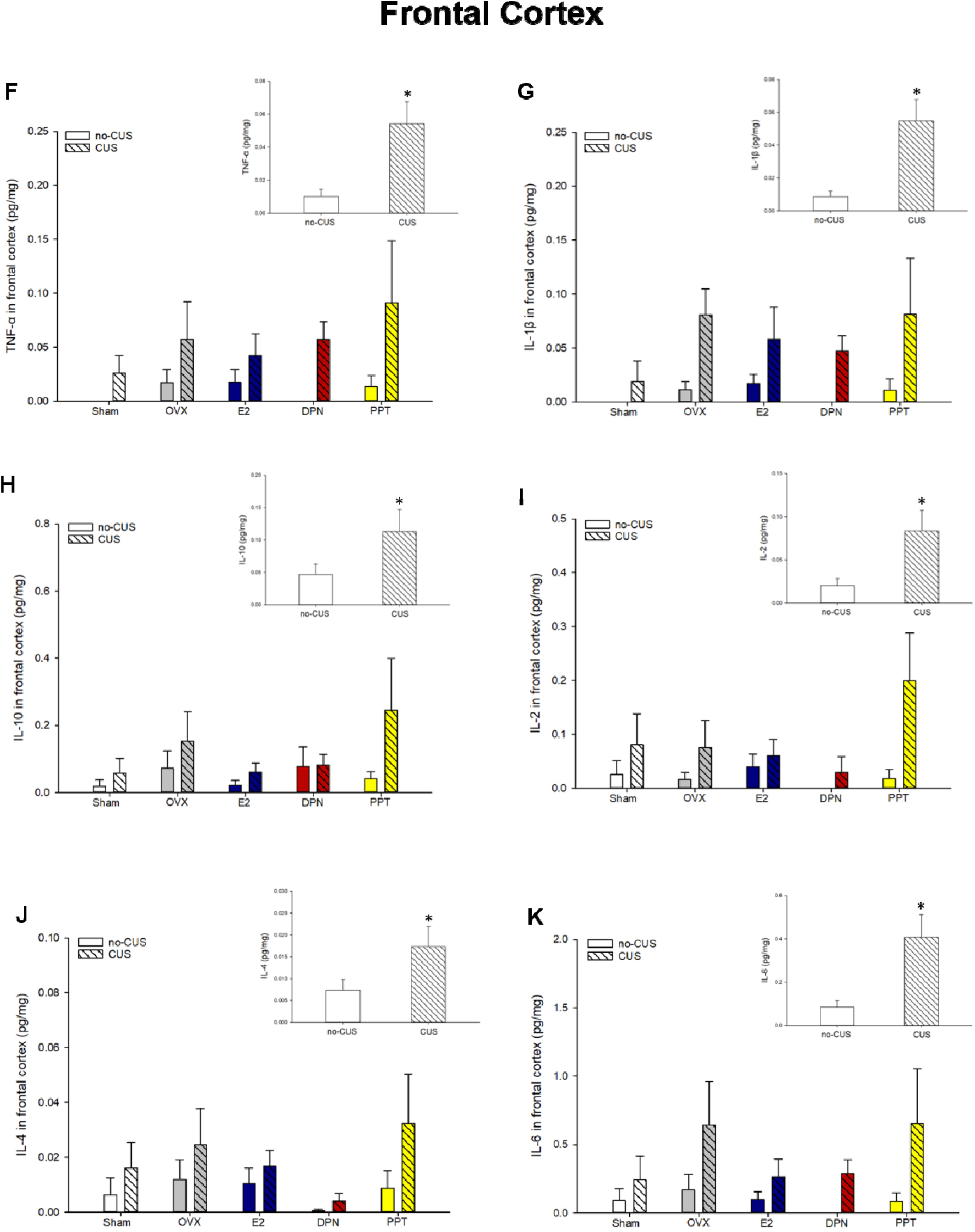
Protein concentrations of immune mediators in the hippocampus (A-E) and frontal cortex (F-K). CUS exposure increased TNF-α **(A)**, IL-1β **(B)**, and IL-10 **(C)** concentrations in the hippocampus; * indicates p <0.015, main effect of CUS condition. **(D)** CUS exposure decreased and E2 treatment increased CXCL1 concentrations in the hippocampus; * indicates p=0.05, main effect of CUS exposure, and # indicates p=0045 relative to Sham. **(E)** DPN treatment significantly increased VEGF concentrations in the hippocampus; * indicates p<0.03, relative to Sham and PPT groups. CUS exposure increased TNF-α **(F)**, IL-1β **(G)**, IL-10 **(H)**, IL-2 **(I)**, IL-4 **(J)**, and IL-6 **(K)** concentrations in the frontal cortex* indicates p ≤0.05, main effect of CUS condition. IL, Interleukin; TNF-α, tumor necrosis factor-α; CXCL1, chemokine (C-X-C motif) ligand 1; VEGF, vascular endothelial growth factor; CUS, chronic unpredictable stress; OVX, ovariectomized; E2, estradiol; DPN, diarylpropionitrile; PPT, propylpyrazole-triol. Data in means + standard error of the mean.

**Table 5.** Protein concentrations (mean ± standard error of the mean) of immune mediators in the hippocampus and frontal cortex, normalised to total protein concentrations (pg/mg). No significant group differences were observed.

|  | Hippocampus |  |  |  |  | Frontal cortex |  |  |  |
| --- | --- | --- | --- | --- | --- | --- | --- | --- | --- |
| Group | GM-CSF | IFN-γ | IL-2 | IL-4 | IL-6 | GM-CSF | IFN-γ | CXCL1 | VEGF |
| **no-CUS** |  |  |  |  |  |  |  |  |  |
| Sham | 0.014 ± 0.0017 | ND | ND | 0.0029 ± 0.0029 | 0.17 ± 0.14 | 0.022 ± 0.005 | ND | 1.37 ± 0.10 | 1.02 ± 0.05 |
| OVX | 0.015 ± 0.0011 | ND | ND | 0.0066 ± 0.0047 | 0.29 ± 0.18 | 0.023 ± 0.006 | ND | 1.99 ± 0.64 | 0.89 ± 0.10 |
| E2 | 0.014 ± 0.0008 | 0.0008 ± 0.0008 | 0.0095 ± 0.0095 | 0.0123 ± 0.0047 | 0.28 ± 0.10 | 0.016 ± 0.005 | 0.0002 ± 0.0002 | 2.18 ± 0.39 | 0.82 ± 0.15 |
| DPN | 0.016 ± 0.0012 | ND | 0.0176 ± 0.0176 | ND | 0.06 ± 0.03 | 0.030 ± 0.006 | ND | 2.34 ± 0.67 | 1.12 ± 0.03 |
| PPT | 0.013 ± 0.0011 | ND | 0.0265 ± 0.0193 | 0.0036 ± 0.0025 | 0.12 ± 0.06 | 0.019 ± 0.004 | ND | 2.33 ± 0.62 | 0.87 ± 0.13 |
| **CUS** |  |  |  |  |  |  |  |  |  |
| Sham | 0.017 ± 0.0024 | ND | 0.0970 ± 0.0706 | 0.0112 ± 0.0051 | 0.19 ± 0.14 | 0.020 ± 0.004 | 0.0008 ± 0.0008 | 1.09 ± 0.09 | 0.82 ± 0.05 |
| OVX | 0.014 ± 0.0015 | ND | 0.0068 ± 0.0068 | 0.0103 ± 0.0048 | 0.50 ± 0.23 | 0.021 ± 0.004 | 0.0002 ± 0.0002 | 1.47 ± 0.28 | 0.94 ± 0.10 |
| E2 | 0.015 ± 0.0025 | ND | 0.0345 ± 0.0227 | 0.0097 ± 0.0055 | 0.25 ± 0.13 | 0.017 ± 0.003 | 0.0002 ± 0.0002 | 2.46 ± 0.79 | 0.95 ± 0.08 |
| DPN | 0.014 ± 0.0022 | 0.0012 ± 0.0008 | 0.0114 ± 0.0114 | 0.0127 ± 0.0089 | 0.34 ± 0.10 | 0.015 ± 0.002 | ND | 1.15 ± 0.13 | 0.81 ± 0.04 |
| PPT | 0.013 ± 0.0024 | 0.0002 ± 0.0002 | ND | 0.0038 ± 0.0038 | 0.13 ± 0.13 | 0.027 ± 0.006 | 0.0044 ± 0.0037 | 1.30 ± 0.08 | 0.87 ± 0.07 |

ND, non-detectable; CUS, chronic unpredictable stress; Sham, sham-operated; OVX, ovariectomized; E2, estradiol; DPN, diethylpropionitrile; PPT, propylpyrazole-triol; IL, Interleukin; IFN-γ , Interferon-gamma; CXCL1, chemokine (C-X-C motif) ligand 1; GM-CSF, Granulocyte-macrophage colony-stimulating factor; VEGF, vascular endothelial growth factor.

In the frontal cortex, CUS exposure significantly increased TNF-α, IL-10, IL-1β, IL-2, IL-4, and IL-6 concentrations (main effects of CUS condition: F(1, 48)=10.59, p=0.0021, η^2^_p_=0.18; F(1,46)=4.00, p=0.05, η^2^_p_=0.08; F(1, 42)=15.63, p=0.0003, η^2^_p_=0.27; F(1, 44)=7.47, p=0.009, η^2^_p_=0.15; F(1, 46)=4.79, p=0.034, η^2^_p_=0.09; and F(1, 44)=9.57, p=0.0034, η^2^_p_=0.18, respectively; **Fig. 7F-K**). However, GM-CSF, IFN-g, VEGF, and CXCL1 concentrations were not significantly affected by CUS exposure (p’s>0.09; **Table 5**). Estrogens did not significantly influence any cytokine in the frontal cortex (all p’s >0.13), nor were there any significant CUS condition by treatment interactions for any cytokine (p’s >0.055). Finally, there was a significant covariate effect of DEX to reduce IL-1β, IL-6, and VGEF in the frontal cortex (p’<0.02), and a trend toward significance for reduced TNF-α (p=0.065), but not for any other cytokine examined (p’s >0.13).

### DPN treatment increased Iba-1 optical density in the dorsal GCL

Irrespective of CUS condition, DPN treatment increased Iba-1 optical density in the dorsal GCL in comparison to OVX and E2-treated mice (p’s <0.05; main effect of treatment: F(4, 53)=2.98, p=0.027, η^2^_p_=0.18; **Fig. 8**). There were no group differences in Iba-1 optical density in the ventral GCL, IL-mPFC, or PL-mPFC (all p’s >0.2; no significant main effects or interactions; **Table 6**).

**Figure 8.**
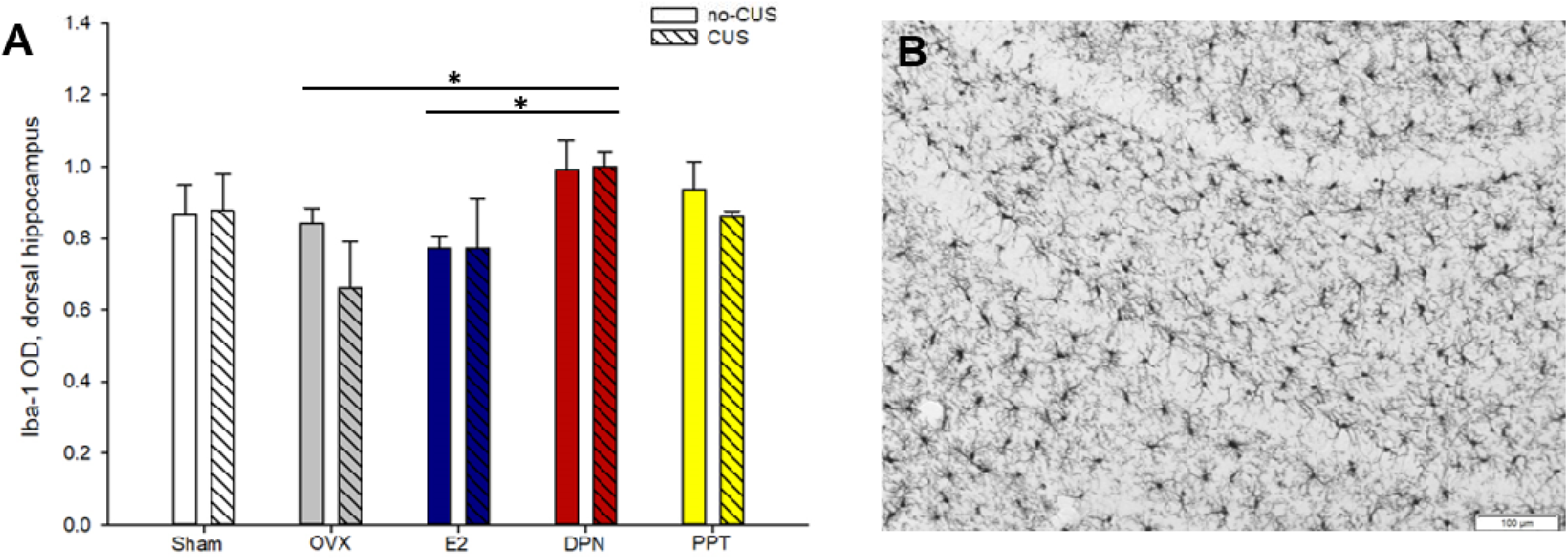
Iba-1 optical density in the dorsal dentate gyrus. **(A)** DPN treatment increased Iba-1 optical density in the granule cell layer of the dorsal dentate gyrus, irrespective of CUS exposure; * indicates p’s <0.05, significantly higher than OVX and E2 groups. **(B)** representative photomicrograph of the dentate gyrus obtained at 10x magnification showing Iba-1-immunoreactive cells. OD, optical density; CUS, chronic unpredictable stress; OVX, ovariectomized; E2, estradiol; DPN, diarylpropionitrile; PPT, propylpyrazole-triol. Data in means + standard error of the mean.

**Table 6.** Iba-1 optical density (mean ± standard error of the mean) in the ventral dentate gyrus and ventromedial prefrontal cortex. No significant group differences were observed.

| Group | vDG | PL vmPFC | IL vmPFC |
| --- | --- | --- | --- |
| **no-CUS** |  |  |  |
| Sham | 0.49 ± 0.02 | 0.74 ± 0.04 | 0.75 ± 0.05 |
| OVX | 0.45 ± 0.02 | 0.71 ± 0.03 | 0.74 ± 0.04 |
| E2 | 0.51 ± 0.03 | 0.74 ± 0.02 | 0.74 ± 0.02 |
| DPN | 0.47 ± 0.05 | 0.75 ± 0.03 | 0.80 ± 0.02 |
| PPT | 0.45 ± 0.04 | 0.71 ± 0.04 | 0.73 ± 0.05 |
| **CUS** |  |  |  |
| Sham | 0.51 ± 0.02 | 0.77 ± 0.02 | 0.78 ± 0.02 |
| OVX | 0.48 ± 0.02 | 0.72 ± 0.03 | 0.69 ± 0.03 |
| E2 | 0.47 ± 0.03 | 0.74 ± 0.03 | 0.77 ± 0.06 |
| DPN | 0.53 ± 0.02 | 0.73 ± 0.04 | 0.76 ± 0.03 |
| PPT | 0.43 ± 0.01 | 0.81 ± 0.06 | 0.81 ± 0.06 |

CUS, chronic unpredictable stress; Sham, sham-operated; OVX, ovariectomized; E2, estradiol; DPN, diethylpropionitrile; PPT, propylpyrazole-triol; vDG, ventral dentate gyrus; vmPFC, ventromedial prefrontal cortex; PL, prelimbic; IL, infralimbic

## Discussion

This study employed a pharmacological approach to investigate the role of the classical estrogen receptors, ERα and ERβ, in the behavioural, neuroplastic and neuroinflammatory consequences of chronic stress exposure in female mice. CUS exposure increased immobility in the TST, increased concentrations of neuroinflammatory mediators, reduced PSD95 expression, and reduced neurogenesis in the ventral hippocampus. The effects of ovarian status and estrogenic treatments were overall less robust than those of CUS. We do, however, observe interesting effects of estrogenic treatments that in some instances depended on CUS exposure. We found that irrespective of CUS exposure, ovarian hormone deprivation increased passive-coping behaviour in the tail suspension test in comparison with sham-operated mice. Intriguingly, E2, DPN, and PPT treatments rescued this depressive-like outcome of ovariectomy under non-CUS conditions, but the effect of CUS exposure to increase depressive-like behaviour was primarily driven by the same treatments. On the other hand, ovarian hormone deprivation produced an anxiolytic effect in the novelty suppressed feeding test regardless of CUS exposure, and this was prevented by estradiol, but not DPN or PPT treatment. Further, the overall effects of CUS to alter the cytokine milieu and to reduce PSD-95 expression in the hippocampus and frontal cortex appear to be driven, at least in part, by DPN and PPT treatments, respectively. Finally, we observed a dorsal hippocampus-restricted effect of estradiol to increase neurogenesis. Combined, these findings shed light on the complexities of estrogen signaling in modulating behavioural, neuroplastic, and neuroinflammatory states under non-stress and chronic stress conditions. These findings further suggest that contrary to our expectations, chronic selective activation of ER α or β does not impart resilience in the face of CUS exposure. These findings could have implications for the development of selective estrogen-based therapies for stress-related disorders.

### Ovarian status and estrogenic treatments produce distinct behavioural profiles depending on CUS condition and behavioural test

Irrespective of CUS condition, ovariectomy increased passive-coping behaviour in the tail suspension test, and under non-CUS conditions, this effect was prevented by E2, DPN, and PPT treatments. This finding is consistent with previous reports showing that ovarian hormone deprivation alone increases depressive-like behaviour in both rats and mice, at least in the short term (Bekku and Yoshimura, 2005; Li et al., 2014). Importantly, our findings implicate both ERα and β in mediating the effects of estradiol to promote active-coping behaviour under non-stress conditions. This counters the findings of past studies that have administered ER agonists acutely to suggest a role of ERβ, but not ERα, in depressive-like behaviour under non-stress conditions (Walf et al., 2004; Yang et al., 2014). This contradictory finding is important, as acute treatments with ER agonists provide only limited information regarding the role of ERs in affective behaviour. Interestingly, although CUS exposure did not significantly affect passive-coping behaviour in the TST in sham-operated and ovariectomized mice, we report that CUS significantly increased passive-coping behaviour in DPN- and PPT-treated mice, with trends in the same direction with E2 treatment. This is somewhat in line with our analysis of resilience versus susceptibility to a depressive-like phenotype in TST, in which CUS exposure resulted in a resilient phenotype in the Sham group but a susceptible phenotype in OVX and PPT-treated groups, while no bias was observed in E2- or DPN-treated groups. This suggests that chronic, selective activation of ERα or ERβ, is protective under non-stress conditions but not under CUS exposure. This seemingly contradictory finding could perhaps be interpreted in light of the “healthy cell bias of estrogen action” hypothesis (Brinton, 2008), put forth to explain observations in which the actions of estrogens result in disparate outcomes in health versus pathology. That is, under “healthy”, non-stress conditions, estradiol and ER agonists protect against ovariectomy-induced depressive-like behaviour, yet under “pathological” chronic stress conditions the same treatments were not protective. A “pathological” state under chronic stress conditions could be related to consequences of increased HPA axis activity, including increased concentrations of corticotropin-releasing hormone (CRH) and corticosterone. This finding is important as it underscores the need for using a model of depression when investigating estrogens’ regulation of affective function.

It is important to note that the effects observed in the TST were not consistent across the entire battery of behavioural tests used in this study. For example, we do not see significant group differences in the forced swim test or the sucrose preference test. It is plausible that testing order could affect behaviour on subsequent tests, an issue that can be addressed by counterbalancing testing order in future studies. Further, unlike the observed effect of ovariectomy to increase passive-coping behaviour in the TST, we found an anxiolytic effect of ovariectomy in the novelty suppressed feeding test. This was seen irrespective of CUS exposure, and was prevented by E2, but not DPN or PPT treatment. This indicates that targeting ERα or ERβ alone was not sufficient to return anxiety-like behaviour to levels similar to that of intact mice. On the other hand, E2 treatment mimicked the phenotype of intact mice, indicating that concurrent activation of both receptor subtypes or possibly the G protein coupled estrogen receptor 1 (GPER) may be involved in this response. In line with our current findings, estradiol benzoate treatment in ovariectomized C57BL/6 or Swiss Webster mice, administered chronically via subcutaneous capsule, increased anxiety-like behaviour in comparison to vehicle treatment across several tests of anxiety-like behaviour (Morgan and Pfaff, 2002, 2001). However, more studies find anxiogenic effects of ovariectomy and anxiolytic effects of acute, sub-chronic, or chronic estradiol treatment in ovariectomized rats or mice (Kastenberger et al., 2012; Walf et al., 2009b; Walf and Frye, 2010). Further, reduced anxiety-like behaviour is observed in proestrus, during which endogenous 17β-estradiol is elevated (Frye et al., 2000; Walf et al., 2009a). Importantly, few studies have investigated the effects of ovarian hormones in the novelty suppressed feeding test, which targets motivated behaviour in addition to anxiety-like behaviour. In middle aged rats, we previously found increased anxiety like-behaviour in NSF with long-term ovariectomy and chronic stress exposure (Mahmoud et al., 2016). Another study in young adult rats found no effect of ovariectomy or chronic estradiol treatment (18 days) on behaviour in NSF (Gogos et al., 2018). It is therefore difficult to synthesize the existing literature, but it appears that length of ovarian hormone deprivation, estradiol dose and treatment regimen, species, strain, age, and test of anxiety-like behaviour can all contribute to the differences in findings.

In this study we investigate the corticosterone response to an acute stressor with and without a dexamethasone challenge (discussed ahead), but we did not explore changes in baseline HPA activity across the experimental timeline. Thus, in order to clarify the complexities and inconsistencies in the literature surrounding the role of ovarian hormones in anxiety- and depressive-like behaviour, it would be important to consider changes CRH and corticosterone as a result of ovariectomy and estrogenic treatments under both non-stress and chronic stress conditions. These hormones are regulated by estrogens (Goel et al., 2014; Oyola and Handa, 2017) and are key effectors of the primary neuroendocrine stress response system with notable effects on anxiety- and depressive-like behaviours, and known perturbations in MDD (Binder and Nemeroff, 2010; Stetler and Miller, 2011).

### Ovarian status and ER agonists influenced the neuroinflammatory consequences of CUS exposure in a brain region-specific manner

The neuroinflammatory effects of chronic stress exposure are well documented in males (Kreisel et al., 2014; Kubera et al., 2011). Here, we also observe robust neuroinflammatory effects of CUS exposure in the frontal cortex and hippocampus in young adult female mice. Notably CUS increased concentrations of IL-1β, IL-6 and TNF-α. This mirrors meta-analyses indicating increased concentrations of the same cytokines in the blood and cerebral spinal fluid of individuals with MDD (Dowlati et al., 2010; Haapakoski et al., 2015; Liu et al., 2012; Wang and Miller, 2018). We also demonstrate that ovarian status and selective ER activation can influence the neuroinflammatory response to stress in a brain region-dependent manner. Specifically, PCA analyses indicate that the neuroinflammatory effects of CUS were driven by PPT-treated groups in the frontal cortex, and by Sham and DPN-treated groups in the hippocampus. This suggests that chronic ERα and ERβ activation potentiated the neuroinflammatory consequences of stress exposure in the frontal cortex and hippocampus, respectively. This regional specificity also agrees with our observed effects of the ERβ agonist DPN to increase Iba-1 optical density in the dorsal dentate gyrus. It is possible that the exaggerated neuroinflammatory response to CUS in DPN and PPT groups could contribute to the enhanced behavioural susceptibility to CUS in the tail suspension test, however the current experiment cannot confirm this link. Previous studies provide support for a role of ERα and ERβ in the anti-inflammatory and neuroprotective actions of estradiol (Brown et al., 2010; Chakrabarti et al., 2014; Lewis et al., 2008; Smith et al., 2011), however these conclusions are largely derived from studies using acute inflammatory challenges *in vivo* or from *in vitro* studies. Therefore, our current findings provide insight into the immunomodulatory actions of estradiol in the context of chronic stress exposure. Although estrogens themselves possess immunomodulatory actions, future studies should also consider how estrogens may influence the neuroinflammatory milieu indirectly via the regulation of the HPA axis, especially under conditions of chronic stress. This is important as CRH can produce anti-inflammatory effects indirectly via the actions of glucocorticoids, but can also can directly produce pro-inflammatory effects (Bellavance and Rivest, 2014; Bereshchenko et al., 2018; Elenkov et al., 1999). Therefore, considering well known HPA-HPG interactions (Goel et al., 2014) may shed light on how differing backgrounds of ovarian hormones can influence the neuroinflammatory consequences of stress exposure.

### 17β-estradiol increased neurogenesis in the dorsal hippocampus

Chronic 17β-estradiol treatment increased the number of BrdU-ir cells in the dorsal dentate gyrus and increased the proportion of BrdU-ir cells that also expressed the mature neuronal marker (NeuN). This finding is interesting considering the functional heterogeneity along the dorsal-ventral axis of the hippocampus, in which the dorsal region is predominantly implicated in cognitive function (Fanselow and Dong, 2010; Strange et al., 2014). Few studies to date have examined the effects of chronic 17β-estradiol treatment in ovariectomized females on the survival of new neurons in the hippocampus, but the available evidence indicates that the effects depend on the timing of 17β-estradiol exposure in relation to that of cell proliferation. The survival of new neurons was found to be increased in cell populations that had proliferated after the initiation of estradiol treatment (McClure et al., 2013), but reduced in cell populations that proliferated prior to the initiation of estradiol treatment (Barker and Galea, 2008; Chan et al., 2014). Our current data corroborate this past literature, as 17β-estradiol treatment here was initiated two weeks prior to BrdU administration. Therefore, our current findings provide further support for the notion that a 17β-estradiol rich vs deficient environment during cell proliferation can largely influence the effects of chronic 17β-estradiol treatment on the survival of new neurons. Further, as several studies report pro-proliferative effects of 17β-estradiol that depend on dose and timing of treatment (Barha et al., 2009; Ormerod and Galea, 2001; Tanapat et al., 2005, 1999), the effect of E2 to increase BrdU-ir cells in the current study could be attributed to increased cell proliferation. To our knowledge, the effects ERα and β agonists on hippocampal cell survival have not been investigated prior to this study. We found that unlike E2 treatment, DPN and PPT did not increase neurogenesis in the dorsal dentate gyrus, despite previous data showing that acute administration of DPN or PPT enhances cell proliferation, albeit at a different dose (Mazzucco et al., 2006). Therefore, promoting the survival of cells proliferating in an estradiol rich environment may require the concurrent engagement of ERα and β or alternatively GPER, which is achieved by estradiol but not PPT or DPN. It is important to note that although NeuN remains to be the gold standard for identifying neuronal fate of adult born granule cells, NeuN immunoreactivity and antibody binding can differ between neurons in other brain regions and is influenced by a variety of factors including protein phosphorylation (Gusel’nikova and Korzhevskiy, 2015; Lind et al., 2005). Thus, the possibility exists that hormonal treatments in this study may have affected NeuN immunoreactivity and colocalization with BrdU without having any significant change in neuronal number. However, in unpublished observations we did not detect any change in NeuN intensity between groups.

### CUS exposure reduced markers of neuroplasticity (PSD-95 expression and neurogenesis)

CUS significantly reduced neurogenesis in the ventral, but not dorsal, hippocampus, and impacted cell fate by reducing the proportion of new cells becoming neurons. This is in line with the finding that neurogenesis in this region promotes stress resilience in males (Anacker et al., 2018) and with an overall role of the ventral hippocampus in stress regulation (Bagot et al., 2015; Fanselow and Dong, 2010; Padilla-Coreano et al., 2016). We also observe an overall effect of CUS to reduce PSD-95 in both the hippocampus and frontal cortex, pointing to a reduction in excitatory synapse number and/or stability. Importantly, this parallels findings of reduced expression of PSD-95, synapse loss, and reduced expression of synapse-related genes in the PFC and hippocampus of individuals with MDD (Duric et al., 2013; Feyissa et al., 2009; Kang et al., 2012). Interestingly, CUS-induced reductions in PSD-95 expression were largely driven by DPN and PPT treated groups. This indicates that chronic DPN or PPT treatment in tandem with CUS exposure may result in remodeling of neuronal circuits in regions important for mood and stress regulation, and this could be linked to the observed behavioural alterations under CUS exposure in the tail suspension test. This effect may be driven by stress-induced alterations in the HPA axis, including possible elevations in basal levels of CRH and corticosterone, a possibility that should be explored in future studies. On the other hand, we found limited effects of estrogenic treatments on PSD-95 expression under non-CUS conditions. In contrast, previous studies report increased PSD-95 expression in the hippocampus of female rats after acute treatment with estradiol, PPT, and to a lesser extent DPN (Waters et al., 2009) and with 17β-estradiol treatment in vitro (Akama and McEwen, 2003), therefore the differences between these findings and our current observations are likely due to treatment duration.

### Chronic 17**β**-estradiol treatment blunted the corticosterone response to an acute stressor and CUS exposure increased HPA axis negative feedback sensitivity

Chronic 17β-estradiol treatment, regardless of CUS exposure, blunted the corticosterone response to an acute restraint stressor. This partially contrasts previous studies which in general have indicated that ovarian hormones, and 17β-estradiol in particular, potentiate acute stress-induced activation of the HPA axis (Carey et al., 1995; Figueiredo et al., 2007; Serova et al., 2010). Importantly, most past studies have used shorter durations of estradiol replacement than in the current study, which likely contributes to the differences in findings. It is important to keep in mind that although the dose of 17β-estradiol used here produce proestrus-like circulating concentrations, the daily treatment regimen does not mimic natural estradiol cyclicity in the mouse, but rather is informative to potential effects of chronic estrogen-based therapies. Our current findings therefore extend the past literature to suggest that longer durations of 17β-estradiol replacement may suppress the HPA axis response to stressors, which could represent a protective effect.

Studies in humans generally report a failure of dexamethasone to suppress cortisol release in a subpopulation of individuals with MDD (Ising et al., 2007; Stetler and Miller, 2011), pointing to impaired HPA axis negative feedback inhibition. In the current study, however, we observed enhanced dexamethasone suppression of corticosterone in mice exposed to CUS, pointing to increased sensitivity of negative feedback inhibition. This effect was present in all treatment groups except for E2, however this could be explained by the above-mentioned suppression of corticosterone in response to acute restraint stress in absence of dexamethasone. This unexpected enhancement of negative feedback inhibition with CUS could be an adaptive response to prevent prolonged exposure to glucocorticoids under chronic stress conditions. Interestingly, a similar phenotype is reported in clinical populations of trauma exposure and post-traumatic stress disorder (PTSD), in individuals with comorbid MDD and PTSD (Griffin et al., 2005; Yehuda et al., 1993), and in women with atypical MDD (Levitan et al., 2002), suggesting that this profile can be seen in certain stress-related psychiatric disorders, including MDD with atypical features, which is more prevalent in women than in men (Marcus et al., 2008).

## Conclusion

In conclusion, we observe depressive-like, neuroinflammatory, and neuroplastic effects of CUS in young adult female mice that were largely independent of ovarian status and estrogenic treatments. However, our findings also point to complex effects of ovariectomy and estradiol/ER agonist replacement that in some instances were dependent on chronic stress condition. Specifically, ovariectomy increased passive-coping behaviour in the tail suspension test regardless of CUS exposure, and this effect was rescued by estradiol and ER agonists only under non-CUS conditions, suggesting that active-coping behaviour is promoted by ERα and ERβ. Intriguingly, the effects of CUS exposure to increase passive-coping behaviour in the TST were largely driven by the same treatments. In agreement with the increased behavioural susceptibility to CUS exposure, the effects of CUS to modify the cytokine milieu and reduced PSD-95 expression appear to be driven in part by DPN and PPT. In addition, although ovariectomy increased passive-coping behaviour in the TST, it reduced anxiety-like behaviour in the NSF test regardless of CUS exposure. This effect was only prevented by 17β-estradiol treatment suggesting that the concurrent activation of ERα, ERβ and GPER produced an anxiogenic effect in this paradigm. We further observe opposing effects of estradiol and CUS exposure on BrdU-ir cell number along the dorsal-ventral axis of the hippocampus, such that 17β-estradiol increased and CUS decreased BrdU-ir cells in the dorsal and ventral regions, respectively. Finally, CUS exposure resulted in increased HPA axis negative feedback sensitivity in most groups, pointing to stress-induced adaptations in HPA axis function. Taken together, these data highlight that estrogenic regulation of behaviour, neuroplasticity, and neuroinflammation is complex and at times depends on chronic stress condition. Moreover, our data indicate that chronic selective activation of ERα or ERβ is not protective under chronic stress exposure, which could have implications for the development of estrogen-based therapies for stress-related disorders such as depression.

